# A Novel Cortico-Striatal NREM Sleep Rhythm in Mice and Non-Human Primates

**DOI:** 10.64898/2026.09.11.750712

**Authors:** Aviv D. Mizrahi-Kliger, Samantha Coury, Cameron L. Woodard, Karunesh Ganguly

**Affiliations:** Neurology Service, San Francisco Veterans Affairs Medical Center, San Francisco, California, USA; Department of Neurology, University of California-San Francisco, San Francisco, California, USA; UCSF-UC Berkeley Graduate Program in Bioengineering, San Francisco, CA, USA

## Abstract

With practice, rapid early gains in performance are followed by a slower phase marked by kinematic refinement, automaticity and enhanced cortical and striatal interactions. While sleep is known to support early learning, its causal role in the slow phase of learning is not known. Here we recorded neuronal activity in primary motor cortex (M1) and the dorsolateral striatum (DLS) during long-term skill acquisition and interleaved sleep. Surprisingly, the slow phase of learning was marked by the emergence of a previously unrecognized 5–10 Hz oscillatory activity during NREM sleep that was coherent across M1 and DLS. This oscillation resulted in repeated joint reactivation of task-specific information in cortex and striatum. Strikingly, during later stages of training, such joint reactivation of task activity increased over the course of NREM sleep, suggesting that sleep-dependent processing strengthens cortico-striatal interactions. The strength of M1–DLS coherence was predictive of next-day performance gains and increased cortico-striatal coupling during task performance. Targeted closed-loop disruption of this oscillation during NREM sleep abolished performance gains, whereas switching to a dose-matched random stimulation paradigm enabled performance improvements in the same animals. Importantly, the same cortico-striatal 5–10 Hz rhythm was also found in sleeping non-human primates, where it was selectively enhanced following learning. Together, we identify, across species, a novel NREM sleep oscillation that is important for sleep-dependent performance gains which depend on cortico-striatal processing.

## Introduction

Sleep is widely implicated in the consolidation of motor skills and cognitive processes^1–3^, yet most studies have focused on short-term or overnight effects^4–9^. Automaticity, the ability to execute motor and cognitive behaviors with minimal effort, is a defining feature of learned behavior and depends on plasticity of cortical inputs to the striatum^10–17^. Increased automaticity, and its role in establishing states of inflexibility, is also implicated in many neurological and psychiatric disorders^18,19^. Because automaticity emerges slowly over prolonged training^16,17,20–24^, it likely reflects long-term changes in cortical and subcortical circuits and may require an interplay between repeated task practice and sleep-dependent processing across days^17,25^. However, the neural mechanisms supporting potential sleep-dependent consolidation across extended timescales remain poorly understood. Here, using dual-site large-scale recordings and closed-loop optogenetics in mice, we tested whether and how sleep-associated neural activity causally shapes the coupling between motor cortex (M1) and the dorsolateral striatum (DLS) during long-term learning. We identified a novel sleep rhythm, termed hereafter cortico-striatal sleep coupling (CSSC), that coordinates sleep-dependent cross-regional reactivation of waking experience and drives plasticity of cortical inputs to the striatum. An analogous cortico-striatal NREM sleep rhythm was also identified in non-human primates, where it was selectively enhanced after learning.

## Results

### A coherent 5–10 Hz cortico-striatal rhythm emerges during skill learning

We trained water-scheduled mice (n=11, n=5 behavior only and n=6 with behavior and electrophysiology) on a reach-to-water task across 6 days of learning, where daily task sessions were followed by sleep sessions (1.5 hours, Fig. 1a). Animals exhibited a consistent learning trajectory where success rate showed marked improvement by day 2 (Fig. 1b–c, day 1 (D1) vs. D2–6 average, 63.94% vs. 80.03%, P = 8 × 10^−4^, paired t-test). Reaction time also exhibited rapid dynamics, with most of the improvement occurring by day 3, although performance continued to improve through day 6 (Fig. 1b, d, D1 vs. D3, 1.71 s vs. 0.54 s, P = 1 × 10^−3^, Wilcoxon signed-rank test, D3 vs. D4-6 average, 0.54 s vs. 0.34 s, P = 9.8 × 10^−3^, Wilcoxon signed-rank test). In contrast, variability in reach trajectory and reach endpoint, quantified as the Euclidean distance between individual reach trajectories and the average trajectory, and the standard deviation of the distribution of individual reach endpoints relative to the average endpoint, respectively, both exhibited a slower gradual improvement from day 1 to day 6 (Fig. 1e–f, normalized trajectory distance: D1–2 average vs. D5–6 average, 1.06 vs. 0.96, P = 6.60 × 10^−3^, paired t-test, normalized reach endpoint standard deviation, D1–2 average vs. D5–6 average, 1.14 vs. 0.86, P = 4.80 × 10^−3^, paired t-test). Indeed, early learning was characterized by higher reaction times, lower success rates and kinematic variability. In contrast, late learning was associated with faster reaction times and with more stereotyped and accurate reaches.

**Fig. 1.**
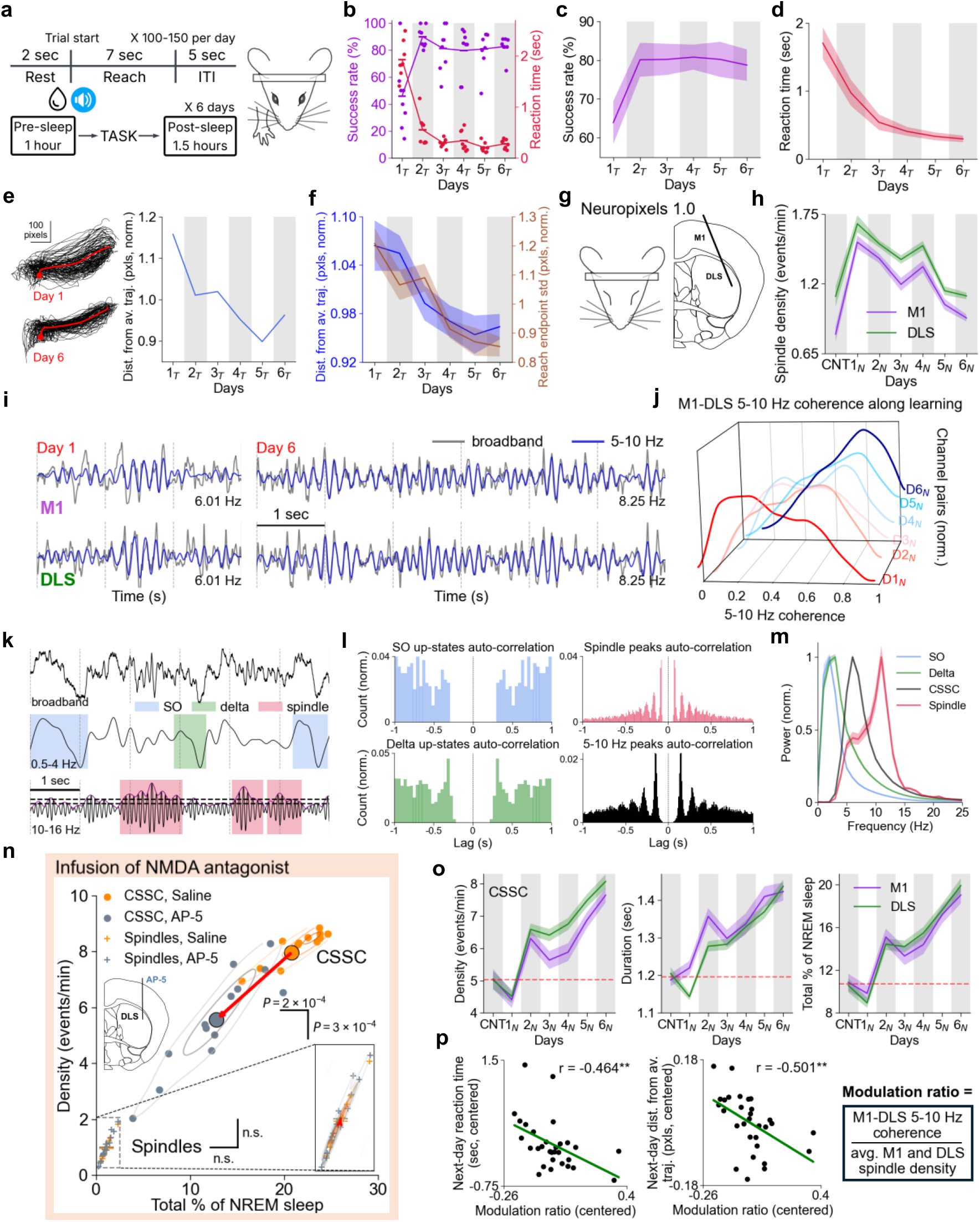
Emergence of a 5–10 Hz sleep oscillation in M1 and DLS that tracks kinematic refinement. **a,** reach-to-water task. Mice reach for a drop of water after an auditory cue, over 6 days. ITI, inter-trial interval. **b,** Success rate (purple) and reaction time (latency from cue to reach onset, red) along days for an example animal. “T” subscript, waking task performance. All per-session trials are grouped into 8 groups of successive trials. Lines connect the session averages. **c,** Success rate for all animals. **d,** Reaction time for all animals. **e,** Left, Reach trajectories for an example animal, with the same number of trials plotted for both days. Red point and trace, average per-day reach start and reach trajectory. Right, Euclidean distance of all reach trajectories from the per-day average reach trajectory, along learning (averaged over x and y coordinates and normalized by the across-day average), same example animal. **f,** Right y axis, average Euclidean distance from average reach trajectory (normalized per animal as above) along learning for all animals. Left y axis, standard deviation of the 2D distribution of single-reach endpoint distances from the per-day average endpoint position (normalized per animal as above) along learning for all animals. **g,** Head-fixed sleep recording immediately after task performance. **h,** Sleep spindle density along days, averaged across channels and across animals. “N” subscript, non-REM sleep (NREMS). CNT, sleep session before task performance on day 1, i.e., prior to the animal’s introduction to the task. **i,** Example M1 and DLS LFP traces during NREMS, early learning (left) and late learning (right), from the same animal. Bottom right, peak frequency of broadband trace (left, middle 1 s. Right, middle 4 s). **j,** Distributions of M1–DLS 5–10 Hz coherence values across all NREMS epochs and animals, along days. **k,** Examples of broadband (0.5–55 Hz) and filtered LFP traces for slow oscillations (SOs), delta oscillations and sleep spindles in M1. Dashed horizontal lines represent the lower and higher instantaneous power thresholds used for detection of sleep spindles (Methods). **l,** Peak auto-correlation diagrams for detected SOs, delta oscillations, sleep spindles and 5–10 Hz oscillation during cortico-striatal sleep coupling (CSSC) events in an example animal. **m,** power spectra of detected SOs, delta oscillations, sleep spindles and CSSC events across all sessions in all animals. Power is normalized to allow direct comparison. Power spectra were obtained from traces 1-s long or longer to allow for a uniform window length (=1 s) and frequency resolution. For SOs and delta oscillations, we used 1-s long segments around the oscillation peak. **n,** Effect of AP-5 infusion into DLS after task training and just prior to sleep on the density and probability of M1 CSSC events and sleep spindles during NREMS. P-values, Mann-Whitney U test. Oval traces represent a Gaussian kernel density estimation. Points and crosses, individual sleep sessions (n=13 for saline, n=14 for AP-5). Larger points and crosses, distribution averages. **o,** Density of CSSC events, duration of individual events and event probability relative to total NREMS duration along days, averaged across channels and animals. Dashed red horizontal lines represent control sleep levels as baseline. **p,** Pearson’s correlation between modulation ratio during sleep and the next-day reaction time (left, P = 9.76 × 10^−3^, Task sessions on days 2-6 with sleep on days 1-5, i.e., five points per animal) or the next-day distance from the average reach trajectory (right, P = 4.75 × 10^−3^), averaged over all trials. All measures are centered by subtracting the across-day mean.

We monitored single-unit activity and local field potentials (LFP) in M1, DLS and mPFC (medial prefrontal cortex) simultaneously using two Neuropixels 1.0 probes during task performance and subsequent sleep (Fig. 1g). We first sought to determine whether these changes in performance corresponded to observable cross-region dynamics during sleep. Consistent with past work^26,27^, we found that post-task non-REM sleep (NREMS) on days 1 and 2 (i.e., the sleep sessions associated with the greatest offline performance gains in success rate and reaction time) was characterized by a significant increase in sleep spindle density (Fig. 1h, M1: control sleep (i.e., pre-task sleep on day 1, before the animal is introduced to the task) vs. D1–2 average, 0.80 vs. 1.46 spindles/min, P = 2.52 × 10^−19^, Wilcoxon signed-rank test, DLS: control sleep vs. D1–2 average, 1.10 vs. 1.59 spindles/min, P = 5.27 × 10^−12^, Wilcoxon signed-rank test). Interestingly, however, from day 3 onwards, sleep spindle density in NREMS exhibited a consistent decrease that persisted until the end of the learning trajectory (M1: D1–2 average vs. D5–6 average, 1.46 vs. 1.02 spindles/min, P = 5.29 × 10^−13^, Wilcoxon signed-rank test, DLS: D1–2 average vs. D5–6 average, 1.59 vs. 1.16 spindles/min, P = 4.24 × 10^−20^, Wilcoxon signed-rank test). Given the inverse relationship between changes in spindle density and the changes over the late learning phase, we assessed if other features of NREMS could account for the differences.

Strikingly, we noticed that M1 and DLS activity during NREMS was associated with episodic bouts of strong 5–10 Hz oscillatory activity that often lasted for several seconds (Fig. 1i). These oscillatory bouts appeared to last longer as the animals became skilled (e.g., days 5-6, when success rate was high and kinematic variability was low), in some instances, lasting up to six seconds (Fig. 1i, S1a). With learning, we also observed an increase in M1–DLS LFP coherence in the 5–10 Hz range during NREMS across animals. We first quantified this by measuring coherence in the 5–10 Hz band across all NREMS, i.e., an unbiased approach that does not classify episodic bouts (D1–2 vs. D5–6, 0.474 vs. 0.623, P = 1.8 × 10^−3^, paired t-test). Across all NREMS and M1–DLS channel pairs, we observed a considerable shift towards higher 5–10 Hz coherence with learning (Fig. 1j, D1 vs. D6, P = 7.56 × 10^−144^, Kolmogorov-Smirnov test). Such an increase can hypothetically reflect a cross-frequency increase in M1–DLS coherence with learning that is not unique to the 5–10 Hz band. To test this, we normalized the 5–10 Hz coherence by the broadband coherence at 2–40 Hz excluding the 5–10 Hz frequency band itself. Normalized 5–10 Hz M1–DLS coherence showed a significant increase with learning across animals (Fig. S1b, D1–2 vs. D5–6, 0.95 vs. 1.04, P = 3.44 × 10^−6^, Wilcoxon signed-rank test). Although strong across M1 and DLS, 5–10 Hz coherence between M1 and mPFC was weak and did not increase with learning (D1–2 vs. D5–6 across animals, 0.318 vs. 0.355, P = 0.151, paired t-test). However, as we have previously shown^20^, learning was accompanied by an increase in M1–mPFC slow oscillation (SO) coupling (Fig. S1c).

Thus, we observed a transition from post-training sleep with relatively elevated spindle density to a regime characterized by higher cortico-striatal coherence in the 5–10 Hz band. We hypothesized that while this regime is present during post-training sleep in the early period, it is markedly enhanced in the late period. To formally quantify 5–10 Hz activity and monitor its evolution along learning, we used the following criteria to detect episodic events (applied to a given M1 or DLS channel): a segment of LFP activity was considered a 5–10 Hz episodic event if it was associated with: (1) 5–10 Hz power higher than the 70th percentile, calculated across all NREMS for that channel, (2) M1–DLS coherence of 0.7 or higher, and (3) duration of 1 second or more (see Methods for data-driven development of criteria). This resulted in the detection of discrete events of high 5–10 Hz power and M1–DLS coherence, which we termed cortico-striatal sleep coupling (CSSC) events. Unlike slow oscillations (SOs), delta waves and sleep spindles, CSSC events had a distinct oscillatory nature that was evident in their waveforms (Fig. 1i, k, S1a), durations (Fig. S1d), auto-correlations (Fig. 1l) and spectral decomposition (Fig. 1m).

Past works in-vitro and under anesthesia have found that convergent cortical inputs to the striatum can trigger “up-states” that are dependent on NMDA receptor activation^28,29^. We thus tested whether striatal NMDA receptor activation is selectively needed for cortico-striatal communication during CSSC. In a separate group of animals that had performed task training, we infused the NMDA receptor antagonist AP-5 (versus saline as a within-animal control) in the DLS and then monitored M1 and DLS activity during NREMS. AP-5 infusion in DLS significantly decreased CSSC event density and probability relative to saline infusion (Fig. 1n, M1 CSSC density, saline vs. AP-5, 7.96 vs. 5.56 events/min, P = 3 × 10^−4^, Mann-Whitney U test, CSSC probability, 20.79% vs. 12.80%, P = 2 × 10^−4^, Mann-Whitney U test). Remarkably, this effect was selective and did not affect sleep spindles (Fig. 1n, M1 sleep spindle density, saline vs. AP-5, 0.96 vs. 0.99 events/min, P = 0.98, Mann-Whitney U test, spindle probability, 1.07% vs. 1.09%, P = 1, Mann-Whitney U test). Thus, unlike sleep spindles, 5–10 Hz activity is dependent on NMDA-mediated activity in DLS. Our results suggest that past in-vitro work on activity-dependent cortico-striatal plasticity might apply to sleep^30,31^.

With learning and across animals, CSSC events became longer and more frequent (Fig. 1o, duration, D1–2 average vs. D5–6 average, M1: 1.29 vs. 1.43 s, P = 2.06 × 10^−12^, Wilcoxon signed-rank test, DLS: 1.21 vs. 1.41 s, P = 4.14 × 10^−20^, Wilcoxon signed-rank test. Density, D1–2 average vs. D5–6 average, M1: 5.36 vs. 7.29 events/min, P = 1.14 × 10^−17^, Wilcoxon signed-rank test, DLS: 5.55 vs. 7.73 events/min, P = 4.45 × 10^−19^, Wilcoxon signed-rank test), accounting for almost 20% of total NREMS by day 6 (ranging between 18.2% to 23.4% across animals, for DLS).

Notably, CSSC event density, duration and probability all exhibited an increase starting on the sleep session immediately after learning day 2: this was the first day on which they exceeded control levels (Fig. 1o, M1, D2 vs. control, density, 6.30 events/min vs. 5.05 events/min, P = 1.11 × 10^−3^, Wilcoxon signed-rank test. Duration, 1.37 s vs. 1.19 s, P = 2.28 × 10^−16^, Wilcoxon signed-rank test. Probability, 15.10% vs. 10.86%, P = 1.70 × 10^−7^, Wilcoxon signed-rank test. DLS, D2 vs. control, density, 6.58 events/min vs. 5.02 events/min, P = 1.99 × 10^−5^, Wilcoxon signed-rank test. Duration, 1.29 s vs. 1.20 s, P = 4.71 × 10^−5^, Wilcoxon signed-rank test. Probability, 14.43% vs. 10.58%, P = 9.46 × 10^−7^, Wilcoxon signed-rank test). This appeared to follow a trajectory similar to the kinematic measures (Fig. 1f). Therefore, to test whether a transition from relatively high spindles to a relatively high CSSC regime could explain changes in behavioral performance, we defined a modulation ratio as the ratio between 5–10 Hz M1–DLS LFP coherence and the average M1 and DLS sleep spindle density, across NREMS. The modulation ratio increased with learning (Fig. S1e, D1–2 average vs. D5–6 average, 0.33 vs. 0.59, P = 2.85 × 10^−3^, paired t-test) and was significantly correlated with next-day reaction time, next-day distance from the average trajectory (Fig. 1p) and next-day reach endpoint standard deviation (Pearson’s r = −0.372, P = 0.042), across animals. Moreover, the next-day distance from the average trajectory was separately positively correlated with M1 and DLS spindle density (i.e., elevated spindle density was associated with more kinematic variability, r = 0.400, P = 0.029) and negatively correlated with 5–10 Hz M1–DLS LFP coherence (r = −0.500, P = 4.85 × 10*^-^*^3^), indicating that kinematic refinement was inversely correlated with spindle activity and positively correlated with 5–10 Hz coherence. Thus, the transition from spindle rich sleep to M1–DLS 5–10 Hz coherent sleep was correlated with early gains in reaction time and late improvement in kinematics.

In some strains of rats, 5–12 Hz activity was reported to associate with whisking/facial twitches during quiet wakefulness or to relate to Absence-like seizure activity^32^. In our animals, CSSC activity was detected during NREMS, and of all CSSC events across animals and sessions, 96.49% were below a mixture of Gaussians model-derived immobility threshold (that was validated to also exclude whisking and facial twitches), i.e., CSSC events were not associated with movements.

In summary, its unique spectral characteristics, apparent abundance in late learning, anatomical confinement to the motor cortical-striatal network, reliance on NMDA transmission and behavioral relevance, collectively establish the 5–10 Hz rhythm as a distinct novel NREMS phenomenon.

### CSSC entrains cross-region spiking and task-specific reactivation

To investigate whether CSSC affects single-neuron spiking, we examined the entrainment of M1 neurons to CSSC events. We hypothesized that as performance becomes more kinematically reliable, spiking in both M1 and DLS will become increasingly entrained to CSSC. We first observed very striking rhythmic modulation of population dynamics in both M1 and DLS in the late period (Fig. 2a shows an example from day 6 NREMS). With learning, M1 and DLS spiking during NREMS became increasingly entrained to CSSC (Fig. 2b–d, M1, all neurons, D1–2 vs. D5–6, phase locking value (PLV) = 0.192 vs. 0.254, P = 4.76 × 10^−35^, Mann-Whitney U test, DLS, PLV = 0.126 vs. 0.159, P = 1.00 × 10^−42^). Further, we noticed that the cross-day distribution of M1 PLVs was bimodal rather than unimodal (Fig. 2b–c, Bayesian Information Criterion minimized for a mixture of n=2 Gaussians); this suggests that the PLV distribution was composed of two distinct distributions, exhibiting low and high CSSC entrainment. Learning was associated with a shift towards higher CSSC entrainment in M1 during NREMS (Fig. 2d). Indeed, we observed the emergence of a population of neurons in M1 exhibiting increased CSSC entrainment (mean PLV = 0.31 across animals). This population accounted for 46.4% of all M1 neurons by day 6, whereas the fraction of neurons belonging to the increased CSSC entrainment population in days 1 and 2 was 18.5% (Fig. 2e, P = 1.73 × 10^−18^, chi-square test).

**Fig. 2.**
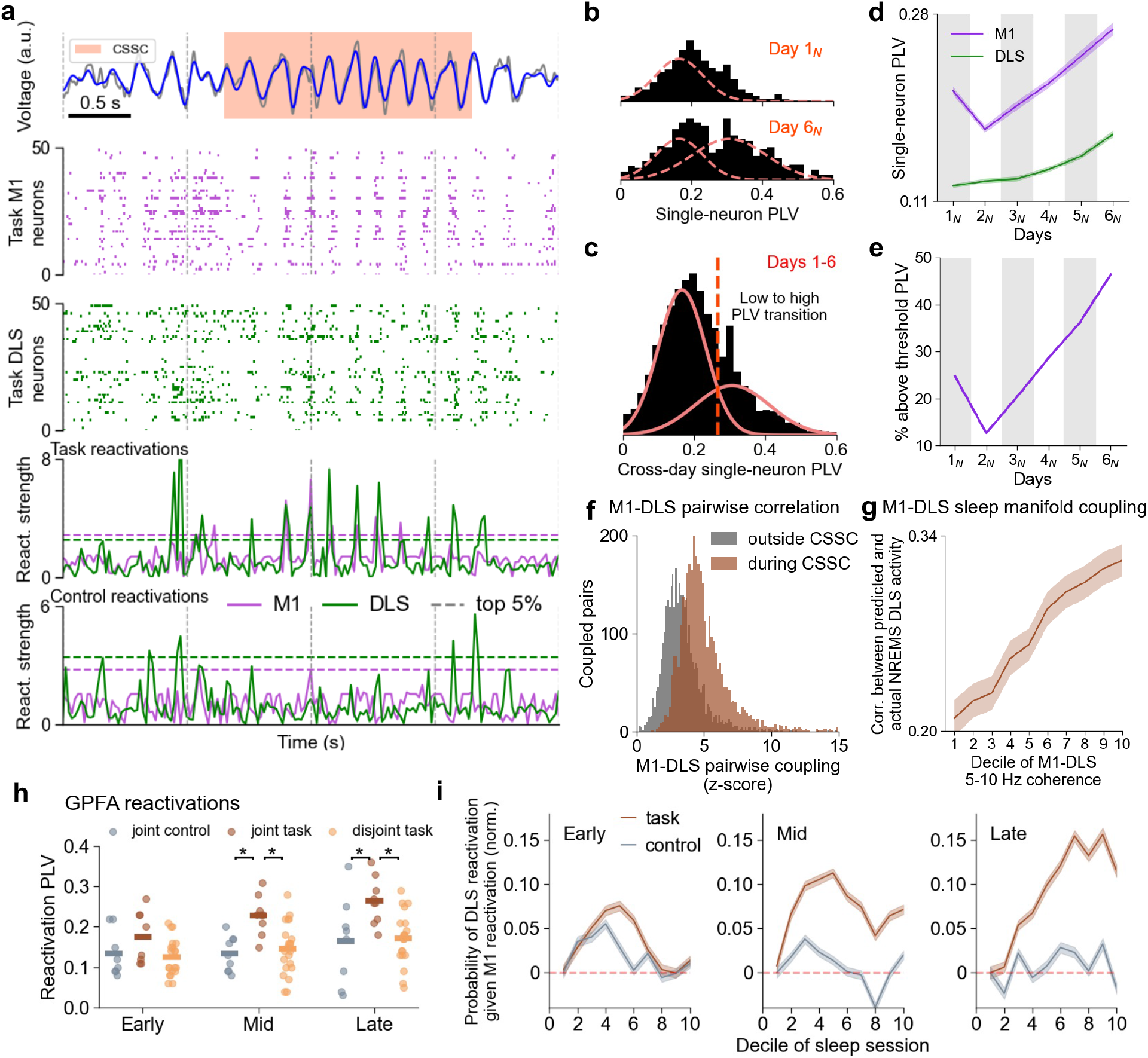
CSSC entrains spiking and task-specific joint reactivation across M1–DLS. **a,** Example 4-s segment showing detected CSSC activity in M1 (top panel. Orange shading, CSSC event, gray, broadband, blue, 5–10 Hz. Dashed vertical lines represent 1 s) along with the activity of 50 representative units in M1 and 50 in DLS (2^nd^ and 3^rd^ panels, respectively, recorded through both task performance and sleep). Bottom two panels, absolute reactivation strength of the first GPFA factor for M1 and DLS. Reactivations are for reach-related information (next-to-bottom panel) or non-reach rest-related information (bottom panel). Horizontal dashed lines represent the top 5% of reactivation strengths across the entire session. **b,** Phase locking values (PLVs) of M1 neurons to the 5–10 Hz oscillation during CSSC events, across all animals, on days 1 and 6 (normalized density). Dashed orange curves represent the two Gaussians underlying the cross-day pooled data (**c**, below), where Gaussian density values were scaled to match per-day histograms for visualization. **c,** Distribution of M1 PLVs during CSSC events, pooled across animals and sleep sessions (n=3,507 neurons). A mixture of Gaussians model revealed that n=2 Gaussian kernels best fit the data (Bayesian Information Criterion of −6012.34 for n=2 in comparison to values ranging from −5589.42 to −5993.40 for n=1, 3, 4). **d,** Average PLVs for all neurons across animals for M1 (n=3,507 neurons) and DLS (n=7,402 neurons). **e,** Percentage of M1 neurons that exhibit PLV values higher than the low to high CSSC entrainment threshold. **f,** Comparison of cofiring of the same M1–DLS neuron pairs during vs. outside of CSSC events (outside, 1-s NREMS segments exhibiting bottom 30% of 5–10 Hz coherence, to allow comparison of a similar duration of time between groups). Pairwise coupling was defined for a pair consisting of a single M1 neuron and a single DLS neuron, as their z-scored cofiring relative to jittered data (Methods). Pooled data from D2-6 (mid + late learning), average z-score during CSSC vs. outside of CSSC, 4.73 vs. 3.26, P = 1.91 × 10^−3^, Wilcoxon signed-rank test. **g,** Pearson’s correlation between actual DLS GPFA trajectories (generated from the top factor) and the trajectories predicted by a Ridge regression model only based on concurrent M1 activity, for 1-s segments during NREMS. Segments are grouped according to the decile of the concurrent M1– DLS 5–10 Hz coherence, such that lower deciles correspond to lower 5–10 Hz coherence. Pooled data from D2-6 (mid + late learning), first two deciles of 5–10 Hz coherence vs. deciles 9–10, r = 0.22 vs. 0.32 across animals, P = 3.42 × 10^−6^, paired t-test. **h,** PLVs of reactivation events to the 5–10 Hz oscillation during CSSC. Joint task, concurrent M1 and DLS reactivations of reach-related information. Disjoint task, separate reactivation events in either M1 or DLS. Joint control, concurrent M1 and DLS reactivations of control information. Joint task vs. joint control, early learning, non-significant, mid learning, P = 1.94 × 10^−3^, and late learning, P = 0.030, Mann-Whitney U test. Joint task vs. disjoint task, early learning, non-significant, mid learning, P = 8.42 × 10^−4^, and late learning, P = 1.11 × 10^−3^, Mann-Whitney U test. Joint task, early vs. late, P = 0.026, disjoint task, P = 0.022, and joint control, non-significant. Horizontal bars, mean PLVs. **i,** Probability of DLS reactivation given M1 reactivation. Reactivation probabilities calculated over deciles of sleep session, across early, mid or late learning sessions, in all animals. Probabilities are normalized by subtraction of the mean probability in decile 1 of the sleep session.

We observed that days 5–6 of the learning trajectory showed more marked increases, relative to earlier days, in M1–DLS coherence (Fig. 1j), CSSC epoch density and duration (Fig. 1o), and M1 and DLS spike locking to CSSC (Fig. 2b–e). Thus, going forward we have grouped our analyses by distinct learning stages across days of the task: early (days 1–2, when spindle densities are high and CSSC is still relatively weak), mid (day 3–4, when CSSC is apparent) and late (days 5– 6).

We then determined whether the entrainment of single neurons in M1 and DLS to CSSC was linked to temporally precise pairwise cofiring of M1 and DLS neurons. To measure functional coupling, we first detected M1–DLS neuron pairs that showed evidence of synchronized firing that is higher than chance for a 40-ms time window across all NREMS (Methods). We found that these coupled cross-region neuron pairs were more likely to fire together during CSSC than outside of CSSC, only for mid and late learning, but not for early learning (Fig. 2f, mid learning, z-scored coupling during vs. outside of CSSC events, averaged across all pairs per session, 4.67 vs. 3.16, P = 9.77 × 10^−4^, Wilcoxon signed-rank test. Late learning, 4.81 vs. 3.39, P = 3.91 × 10^−3^, Wilcoxon signed-rank test. Early learning, 3.88 vs. 3.38, P = 0.083, Wilcoxon signed-rank test. Across animals and sessions, n=5,784 coupled M1–DLS neuron pairs of a total of n=690,270 neuron pairs recorded together). To determine whether this increase in M1–DLS coupling was unique to CSSC events we compared coupling strengths for the same pairs of coupled neurons during CSSC vs. during periods of generally increased broadband coherence. Here as well, we observed stronger M1–DLS coupling during CSSC than during periods of high non-specific broadband coherence (NREMS 1-s segments exhibiting coherence values in the top 30% within the 2–40 Hz range, excluding 5–10 Hz. Mid learning, z-scored coupling during CSSC events vs. broadband, 4.67 vs. 3.89, P = 9.77 × 10^−4^, Wilcoxon signed-rank test. Late learning, 4.81 vs. 3.92, P = 3.91 × 10^−3^, Wilcoxon signed-rank test). This suggests that CSSC not only entrains M1 and DLS firing but is associated with increased pairwise cross-region coupling.

The entrainment of large neuronal populations in M1 and DLS to CSSC as well as the increase in cross-region firing associated with it raises an intriguing possibility that cross-region spiking during CSSC might be instrumental in coupling manifold-level activity across M1 and DLS, not only single neurons. If that were true, the degree of M1–DLS coupling could be predicted based on moment-by-moment CSSC range coherence levels. To test this possibility, we examined M1– DLS coupling during NREMS by monitoring the ability of M1 population activity to predict current DLS population-level spiking. We first used Gaussian-Process Factor Analysis (GPFA^33^) to extract low-dimensional patterns of M1 and DLS activity during NREMS. We then trained a linear regression model to predict bin-by-bin DLS GPFA single-factor activity, based solely on M1 factor activity. Higher population-level coupling would be reflected in greater similarity between the model’s prediction and actual DLS activity.

Indeed, with increasing 5–10 Hz coherence, we observed an increase in the correlation between the decoder prediction and actual DLS activity. Here as well, this was true for mid and late learning, but not for early learning, when CSSC is low (Fig. 2g, mid learning, first two deciles of 5–10 Hz coherence vs. deciles 9–10, Pearson’s correlation between predicted and actual DLS activity, 0.21 vs. 0.29 across animals, P = 1.20 × 10^−3^, paired t-test. Late learning, 0.22 vs. 0.35, P = 6.42 × 10^−4^, paired t-test. Early learning, 0.31 vs. 0.37, P = 0.115, paired t-test). Further, the increase in M1–DLS population level coupling was specific to the 5–10 Hz band: when we examined the performance of the decoder relative to concurrent broadband coherence (i.e., 2–40 Hz, excluding the 5–10 Hz band), we found no difference in correlation between low and high coherence states (mid learning, first two deciles of broadband coherence vs. deciles 9–10, Pearson’s correlation between predicted and actual DLS activity, 0.23 vs. 0.28 across animals, P = 0.230, paired t-test. Late learning, 0.30 vs. 0.32 across animals, P = 0.592, paired t-test).

We next sought to determine if this population-level coupling during sleep could subserve offline memory consolidation underlying kinematic refinement (Fig. 1). To examine this hypothesis, we first tested whether CSSC was associated with coincident cross-region reactivation of task-related information in M1 and DLS, i.e., joint reactivations. We again utilized GPFA to extract low-dimensional patterns of neuronal firing separately in M1 and DLS, specifically focusing on reaching during task performance. Recording the activity of the same M1 and DLS neurons during subsequent sleep, we detected reactivation events of reach-related cofiring patterns by projecting binned activity during NREMS onto the reach-derived factor subspace^34^. Finally, we defined joint M1–DLS reactivations as above-threshold reactivations in both M1 and DLS that occurred in the same time bin (Methods). In addition to reactivations of task-specific activity, we also detected reactivations of waking information that was not associated with reaching, i.e., reactivation of neuronal activity recorded during the enforced rest period before trial start (when animals must keep their paws still to trigger the next trial).

Across animals, M1–DLS joint reactivation density was significantly higher during vs. outside of CSSC events. This was true for all stages of learning, but the difference increased as learning progressed (Fig. S2, during vs. outside, Early learning, 0.85 vs. 0.69 events/s, P = 3.22 × 10^−2^, Wilcoxon signed-rank test. Mid learning, 0.89 vs. 0.67 events/s, P = 4.88 × 10^−3^, Wilcoxon signed-rank test. Late learning, 1.01 vs. 0.68 events/s, P = 4.88 × 10^−3^, Wilcoxon signed-rank test). Specifically, by days 5 and 6, we observed numerous instances where CSSC was associated with repeated reactivation of task-related activity in both M1 and DLS. Remarkably, such activity during late learning could be specifically associated with a high-density coordinated reactivation of task-related, but not control, information (Fig. 2a, bottom panels, and see below for quantification).

Reactivations were both content- and learning stage-specific. Joint M1–DLS reactivations showed a clear phase preference for the CSSC 5–10 Hz oscillation, with reactivations of task information exhibiting stronger phase locking than either joint M1–DLS control reactivations or disjoint task reactivations in M1 or DLS (i.e. occurring in either region separately). These effects emerged during mid and late learning, but not during early learning (Fig. 2h). Moreover, phase locking of joint reactivations to the CSSC 5–10 Hz oscillation strengthened progressively with learning (Fig. 2h, early vs. late sessions PLV = 0.175 vs. 0.264, P = 0.026, Mann-Whitney U test).

What is the functional significance of joint M1–DLS reactivation for memory consolidation? If reactivations were to bias cross-region synaptic weights to induce a persistent change in cross-region coupling, this would be reflected in a change in the ability of M1 reactivations to elicit DLS reactivations along the sleep session. Indeed, we observed a gradual increase in the probability of DLS reactivation given an M1 reactivation along the sleep session. This gradual increase was absent during early learning (first two deciles of the sleep session vs. last two deciles, 0.015 vs. 0.007 increase in probability relative to baseline, P = 0.133, Mann-Whitney U test), appeared during mid learning and peaked during late learning, when CSSC activity and spike and reactivation locking was at its highest (first two vs. last two deciles, −0.0001 vs. 0.135 increase in probability relative to baseline, P = 1.34 × 10^−88^, Mann-Whitney U test). Remarkably, this along-session increase in M1 ability to recruit DLS reactivation was specific to task-related information and not control reactivations (Fig. 2i). The magnitude of the per-decile increases in probability of DLS reactivation given M1 reactivation was correlated with the per-decile probability of CSSC events for late learning (Pearson’s r = 0.351, P = 3.01 × 10^−4^) but not for early or mid-learning (0.139 and 0.120, both non-significant).

Finally, we also analyzed whether joint M1–DLS reactivations might also track spindling between M1 and DLS. Interestingly, joint reactivation density was significantly lower during M1-detected sleep spindles than during CSSC events (across all learning stages, 0.916 events/s vs. 0.278 events/s, P = 1.31 × 10^−10^, Wilcoxon signed-rank test). In addition, unlike CSSC activity, the increase in the ability of M1 reactivations to recruit DLS reactivations was not correlated with either sleep spindle probability or density in any of the stages of learning.

### M1–DLS coupling during awake performance increases with learning and is predicted by CSSC

The above results suggest that CSSC entrains cross-region spiking and joint reactivation events in M1 and DLS in a content-specific manner. What are the possible functional consequences on awake task performance? If joint reactivations facilitate binding of task-specific representations in M1 and DLS, then we would expect that M1 and DLS activity during task performance will become more coupled. Indeed, task performance during early learning appeared to be associated with noisy M1 spiking and a relatively weak tuning of DLS firing to reach onset (Fig. 3a, left). By day 6, however, we observed that skilled behavior was associated with increased coupling of M1 and DLS population firing during reaches, with DLS appearing to show stronger reach-related firing (Fig. 3a, right). Across animals, there was a significant increase in reach-related M1–DLS pairwise coupling with learning (D1 vs. D6, 0.42% vs. 2.91% coupled pairs, P < 10^−300^, chi-square test), whereas control-related increase in pairwise coupling was less pronounced (Fig. 3b, D1 vs. D6, 0.06% vs. 0.25% coupled pairs, P = 1.66 × 10^−29^, chi-square test).

**Fig. 3.**
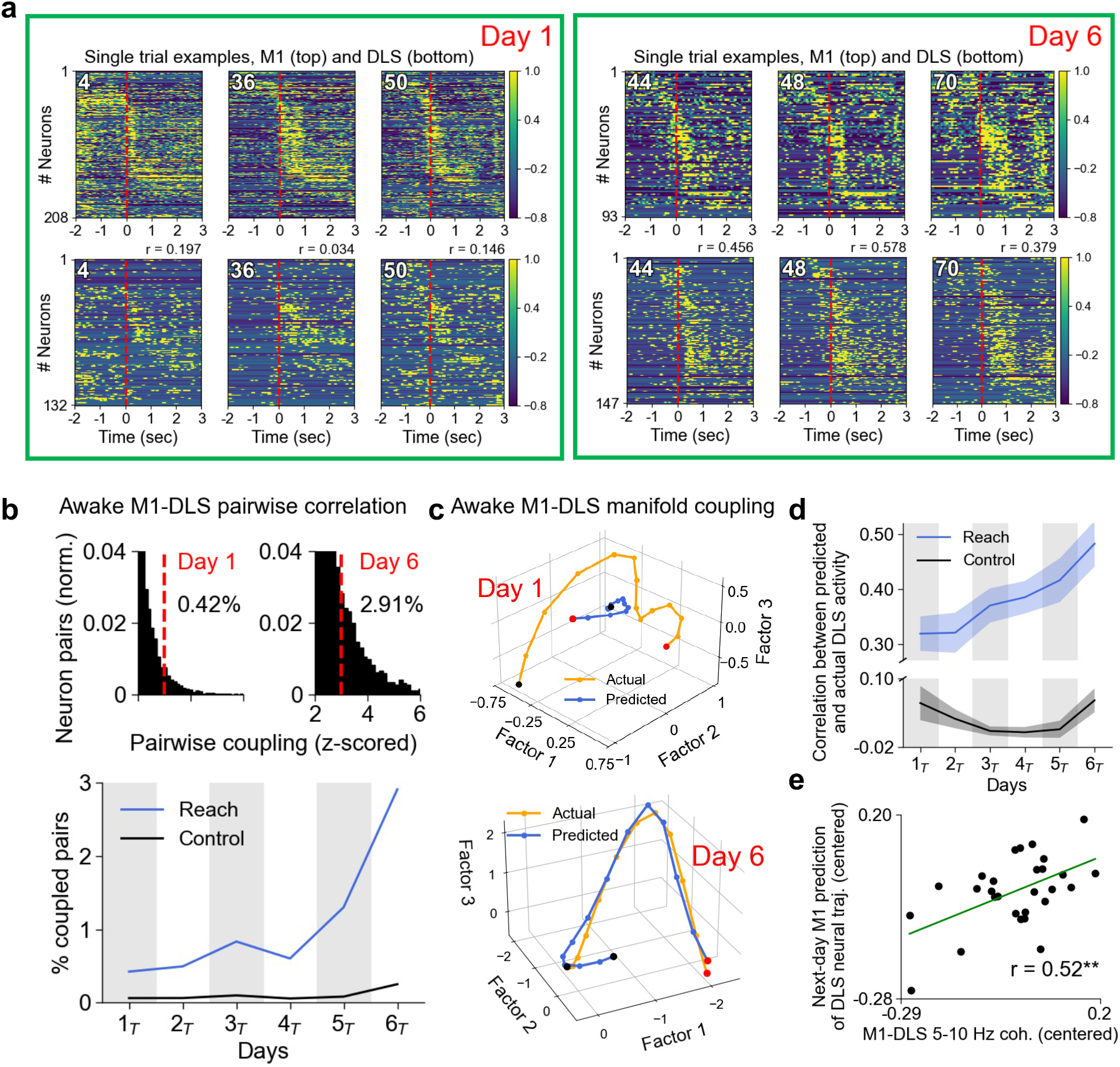
M1–DLS awake coupling tracks learning and correlates with CSSC during previous sleep. **a,** Traces of simultaneously recorded M1 and DLS neurons around reach onset for three example trials in early (Day 1) and late learning (Day 6) from the same animal. Neuronal tiling is based on the time where maximum firing was attained, calculated over all odd trials. Then the same order was applied to even trials (plotted) to test the fidelity of neuronal tuning. Red dashed line, reach onset. Trial numbers on the top left of each panel. Pearson’s r values refer to the correlation between the actual and predicted DLS GPFA trajectories, based on M1 activity (see **c**–**d** below). Color bar scales represent z-scored firing rates. **b,** Top, the proportion of M1–DLS pairs exhibiting significant pairwise coupling (z-scored simultaneous spike count > 3 relative to jittered distribution, Methods) out of all pairs and all animals, for days 1 and 6. Coupling is calculated on a period from 240 ms before to 160 ms after reach onset (to capture immediate preparatory activity as well as the reach itself). Bottom, proportion of coupled pairs, of all simultaneously recorded pairs across days (n=709,628 M1–DLS pairs in total). **c,** Example single-trial GPFA trajectories for predicted (based on concurrent M1 activity) and actual DLS activity during reaching, days 1 and 6. Black and red points indicate trajectory start and end, respectively. **d,** Pearson’s correlation between actual DLS GPFA trajectories and the trajectories predicted by a Ridge regression model (Methods) only based on concurrent M1 activity, during reaches and during a control resting period, across all animals and along learning (averaged over the first 3 factors). **e,** Pearson’s correlation between the average M1–DLS 5–10 Hz coherence during a sleep session and the next-day ability to decode DLS activity based on M1 firing (quantified as the correlation between the predicted and actual GPFA traces). All measures are centered by subtracting the across-day mean.

Given our finding that NREMS CSSC activity was associated with manifold-level coordination of M1 and DLS neural trajectories, we investigated whether population-level changes also occur during awake behavior. We trained a linear regression model on M1 single neuron firing during task performance and examined how well it could predict ongoing DLS neural GPFA trajectories. Indeed, from day 1 to day 6 the model showed significant improvement in prediction of DLS reach activity based on M1 reach activity, while no such trend appeared for a model trained on control period activity (Fig. 3c–d. Reach, Pearson’s correlation between the predicted and actual DLS GPFA trajectory, across animals, D1 vs. D6, 0.33 vs. 0.48, P = 1.16 × 10^−3^, Wilcoxon signed-rank test. Control, D1 vs. D6, 0.08 vs. 0.06, P = 0.454, Wilcoxon signed-rank test). This suggests that manifold-level activity in these two areas became more coupled with learning. Importantly, both M1–DLS 5–10 Hz coherence and CSSC probability in the sleep session prior to task performance were correlated with the next-day performance of the decoder for reaching (Fig. 3e, M1–DLS 5– 10 Hz coherence, Pearson’s r = 0.52, P = 4.27 × 10^−3^. CSSC probability, r = 0.48, P = 0.010) but not for control (M1–DLS 5–10 Hz coherence, r = 0.11, P = 0.580. CSSC probability, r = 0.08, P = 0.681), indicating that offline CSSC predicts next-day neuronal coupling between M1 and DLS during behaviors which are undergoing active kinematic refinement and are associated with successful task behaviors.

### Closed-loop inhibition of DLS during CSSC abolishes offline gains in kinematic refinement

The above results suggest that CSSC facilitates joint reactivation of task-related activity in M1 and DLS and thereby possibly enables kinematic refinement. We directly tested this hypothesis by inhibiting DLS activity specifically during 5–10 Hz oscillations in NREMS (measured in DLS) and assessing how this affected next-day task performance. In a separate experiment, animals (n=6) were implanted with an optrode (32-channel array combined with a fiber-optic cannula, see Methods) following viral infusion of the inhibitory opsin hSyn-eNpHR 3.0 in the DLS. This allowed for simultaneous recording of LFP and spiking as well as concurrent inhibition of the DLS (Fig. 4a). These animals were trained on the reach-to-water task exactly as the control cohort (Fig. 1).

**Fig. 4.**
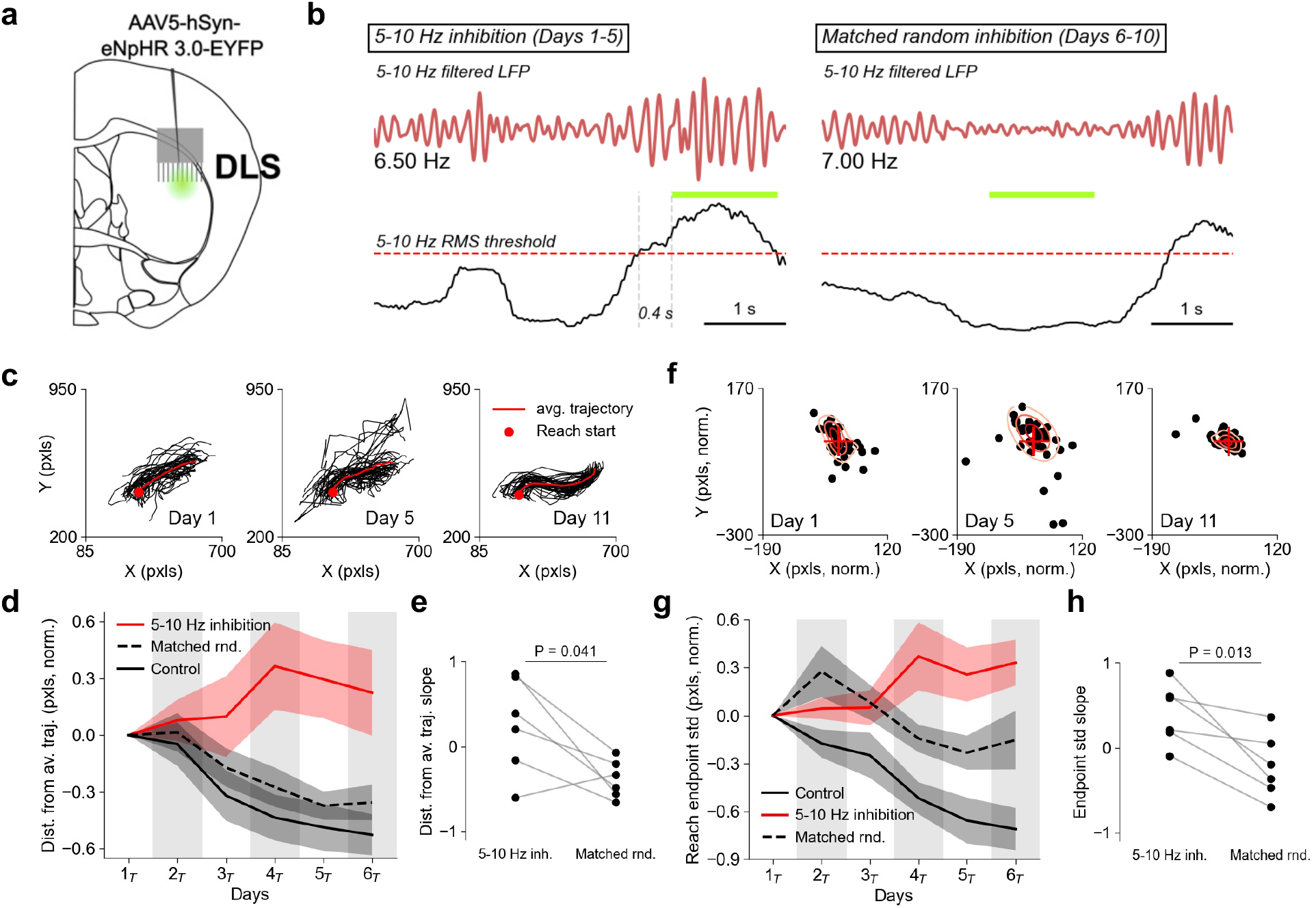
Closed-loop inhibition of DLS during the 5–10 Hz oscillation abolishes kinematic refinement. **a,** Wild-type mice were injected with a viral vector carrying the inhibitory opsin eNpHR 3.0 in the DLS. Later, an optrode consisting of an electrode array and an optic cannula was implanted in the DLS to allow simultaneous recording and optogenetic inhibition. **b,** Closed-loop inhibition policy. In days 1–5 of the intervention, a 532-nm wavelength green-yellow laser was activated after the 5–10 Hz RMS has reached a predefined threshold and stayed above it for 0.3-0.4 s. the laser remained active until the RMS fell below the threshold. In the matched random inhibition regime (days 6–10) the laser was activated randomly during NREMS. The overall duration of laser activation and the average duration of individual laser activations in days 1–5 were used in the matched random regime. **c,** Reach trajectories from day 1, day 5 (last day of 5–10 Hz inhibition) and day 11 (after the matched random inhibition) from an example animal. Trials were randomly subsampled so that the same number of trials is plotted for each day. **d,** Average Euclidean distance of all reach trajectories from the per-session average reach trajectory, along days, for the 5–10 Hz inhibition regime, the matched random inhibition regime and control animals (no optogenetic intervention). All traces are smoothed and normalized by intervention day 1 (day 1 for control and 5–10 Hz inhibition, day 6 for matched random). **e,** Per-animal learning slopes for the change in Euclidean distance from per-session average reach trajectory. For the 5–10 Hz inhibition regime, average of days 5–7 minus day 1. For the matched random, day 11 minus the average of days 5–7. The average is used to account for between-animal differences in behavior dynamics. **f–h,** As **c–e,** but for reach endpoint standard deviation. In **f**, pixel distances are normalized so that the spout location is at the origin. Oval traces represent a Gaussian kernel density estimation.

During post-task sleep in days 1 to 5, we inhibited DLS activity: 5–10 Hz LFP activity was monitored online during NREMS, and when the RMS of this oscillation exceeded a threshold for more than a predefined duration, we activated a laser to inhibit DLS spiking until the 5–10 Hz activity RMS went below the above threshold (“5–10 Hz inhibition” regime, Fig. 4b, left). After 5 days of the above intervention, we employed a “matched random inhibition” intervention for NREMS after days 6–10 of learning. Here, the same overall duration of inhibition as in days 1–5 (in packets of the same average laser activation duration) was used during NREMS, but delivery was unrelated to the 5–10 Hz oscillation (Fig. 4b, right). DLS inhibition was effective in lowering firing rates for putative medium spiny neurons in the DLS by ∼60% (Fig. S3a). In post-hoc detection of putative CSSC events, the laser was ON during 67.4% of the total duration of CSSC events across animals and sessions, and of all laser stimulation duration, 72.2% corresponded to a post-hoc detected putative CSSC event. The average duration of post-hoc detected CSSC events during the 5–10 Hz inhibition regime was not significantly different than their average duration during matched random inhibition (2.02 s vs. 1.85 s, P = 0.075, Mann-Whitney U test). This may be consistent with prior work showing that partial reductions in cortical spiking do not necessarily affect global LFP phenomena^35^.

We then analyzed next-day task performance for the 5–10 Hz inhibition (i.e., days 1 to 6 of learning) and the matched random (days 6 to 11) regimes. We found that kinematic learning in the reach-to-water task was abolished during the 5–10 Hz inhibition period. Rather than showing continuous improvement as in our non-perturbed controls, animals exhibited greatly reduced stereotypy in reach trajectory (Fig. 4c–d, average Euclidean distance from session-average reach trajectory, for days 2 to 6 (D2-6), normalized by D1, 5–10 Hz inhibition vs. control, 0.21 vs. −0.36, P = 7.03 × 10^−6^, Mann-Whitney U test) and reach endpoint (Fig. 4f–g, average standard deviation from session-average reach endpoint, for D2-6, normalized by D1, 5–10 Hz inhibition vs. control, 0.21 vs. −0.46, P = 4.32 × 10^−8^, Mann-Whitney U test).

Strikingly, as soon as we switched to the matched random inhibition, where the same overall duration of DLS inhibition was used, but unrelated to 5–10 Hz activity, the animals showed a rapid improvement in kinematic performance. The slope of cross-day change in variability was positive during the 5–10 Hz inhibition regime for both kinematic measures, indicating a deficit of kinematic refinement (slope of distance from session-average reach trajectory, 5–10 Hz inhibition vs. control, 0.25 vs. −0.53, P = 0.014, Mann-Whitney U test. Slope of reach endpoint standard deviation, 0.40 vs. −0.71, P = 3.01 × 10^−3^, Mann-Whitney U test. See Fig. 4e for calculation of slopes). After switching to the matched random regime, the slope for both measures became negative, indicating between-day improvement and kinematic refinement (Fig. 4e, slope of distance from session-average reach trajectory, 5–10 Hz inhibition vs. matched random, 0.25 vs. −0.38, P = 0.041, paired t-test. Fig. 4h, slope of reach endpoint standard deviation, 0.40 vs. −0.22, P = 0.013, paired t-test). Further, the profile of improvement in trajectory stereotypy for the matched random inhibition period was indistinguishable from normal unperturbed controls (Fig. 4d, average Euclidean distance from session-average reach trajectory, for D2-6, normalized by D1, matched random inhibition vs. control, −0.23 vs. −0.36, P = 0.103, Mann-Whitney U test. Slope for matched random vs. control, −0.38 vs. −0.53, P = 0.301, Mann-Whitney U test). For the reach endpoint variability as well, the matched random inhibition regime was associated with an improvement in performance (Fig. 4g, average standard deviation from session-average reach endpoint, intervention D2–3 average vs. D5–6 average, 0.18 vs. −0.19, P = 1.19 × 10^−3^, paired t-test), similar to control animals but unlike the 5–10 Hz inhibition regime. Importantly, we observed the same success rates for both the 5–10 Hz inhibition regime and the matched random regime (Fig. S3b, 84.11% vs. 86.08%, P = 0.267, paired t-test), indicating that the difference in kinematic performance is not due to a general lack of motivation or an inability to perform the task.

Notably, closed-loop 5–10 Hz inhibition also attenuated the reduction in reaction time seen in early learning (average reaction time, D2-6 relative to D1, 5–10 Hz inhibition vs. control, −0.836 s vs. −1.197 s, P = 2.76 × 10^−2^, Mann-Whitney U test). Unlike the kinematic measures, however, reaction time did not improve further after we switched to the matched random inhibition regime (average reaction time, D2-6 of 5–10 Hz inhibition vs. D2-6 of matched random inhibition, relative to D1, −0.836 s vs. −1.023 s, P = 0.267, Mann-Whitney U test). Notably, secondary motor cortex (M2) and M1 both project to DLS. M2-DLS projections may support preparation, movement onset and reaction time^36–38^. Our observation may further support the possibility that both M2-DLS and M1-DLS projections are modulated by CSSC. Moreover, it is possible that early-stage stabilization of reaction time reflects strategies that are established rapidly during initial execution and remain static even after the 5–10 Hz oscillatory coupling is restored.

In summary, by specifically inhibiting DLS activity during detected epochs of strong 5–10 Hz oscillations in NREMS, these results causally demonstrate that offline 5–10 Hz activity in the DLS is required for kinematic improvement in the reach-to-water task.

### CSSC in non-human primates and reactivity to learning

Our results from mice identify CSSC in M1 and DLS during sleep as a driver of kinematic refinement, presumably by coordinating cross-region reactivation of waking experience. Seeing that this type of activity is enriched in sleep during later learning stages, accounting for 20% of all NREMS, and appreciating that core mechanisms for memory consolidation are shared across species^2,3^, we also aimed to determine if CSSC could also be detected in non-human primates (NHPs). To that end, we re-analyzed LFP recordings from the putamen and frontal epidural ECoG of NHPs sleeping after performing a cognitive-motor task (Fig. 5a–b). Before sleep, food-scheduled primates performed a task in which three fractal cues predicted positive (juice reward), negative (air puff), or neutral outcomes. In each block, animals were presented with 10 trials of each cue type, for a total of 30 trials, across 3–4 blocks before sleep. Individual blocks consisted of either familiar or novel cues (i.e., cues that the animals had not seen before). For the novel cues (1–2 each pre-sleep session), the primates quickly learned to anticipate the outcomes, licking in response to reward-predicting cues and blinking in response to cues predicting air-puffs^39^.

**Fig. 5.**
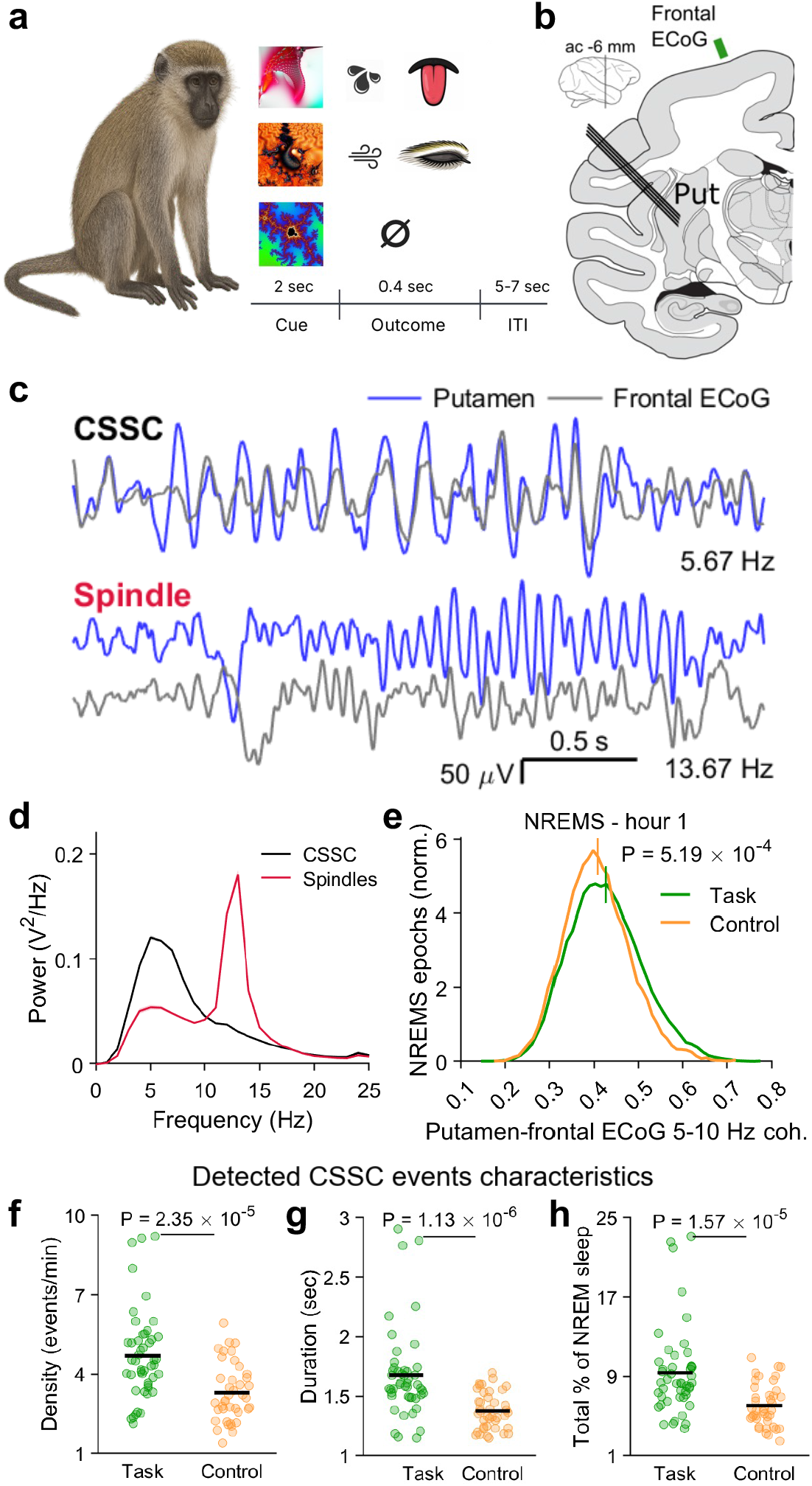
CSSC in NHPs and reactivity to learning. **a,** Green African monkeys performed a classical conditioning task prior to sleep, where they learned to associate fractal cues with rewarding, aversive or neutral outcomes. Within-block learning of new associations resulted in anticipatory licking and or blinking after several cue presentations. Eye and tongue illustrations were generated with AI. **b,** Frontal (as well as central and occipital) ECoG activity was recorded during NREMS, along with neuronal activity in the left putamen. ac, anterior commissure. put, putamen. **c,** Example of 5–10 Hz activity in both the putamen and the frontal ECoG (top), and sleep spindles in the putamen and ECoG (bottom). Broadband filtration at 3-30 Hz. Both ECoG traces were amplified by 10% and ECoG spindle trace was offset for clearer visualization. **d,** Power spectra of detected sleep spindles and CSSC events in the putamen, across all electrodes and sessions in both animals. **e,** Unbiased putamen-frontal ECoG coherence in the 5–10 Hz range, in the first hour of sleep after task performance, vs. in the first hour of sleep when no task has been performed. Vertical bars, average. P-value, Mann-Whitney U test. **f–h,** CSSC event density, duration and overall probability across putamen electrodes and sessions, task vs. control. P-values, Mann-Whitney U test.

Importantly, unlike the mice, the NHPs were extensively trained for months on understanding the cognitive aspects of the task; then each day they had to relearn the association between new fractal cues and a familiar set of a positive, negative and neutral outcomes. As outlined in the discussion, this task may be analogous to rapid sequence learning in human subjects^40^ and in primates^41^.

Recording from the putamen (analogous to rodent DLS) and from the frontal ECoG^42^ approximately above supplementary motor area (SMA) to pre-SMA border (analogous to rodent M1–M2, see Methods), we found evidence for strong 5–10 Hz activity in NHPs during NREMS after task performance (Fig. 5c). As with our findings in mice, this oscillatory activity was different than sleep spindles in both appearance and spectral content (Fig. 5c–d). Using an unbiased approach that does not classify episodic bouts, we compared putamen-frontal ECoG coherence between NREMS after a preceding task training and control sleep (where no task training has occurred before sleep). We found that post-task NREMS was associated with a significant increase in putamen-frontal ECoG 5–10 Hz coherence (Fig. 5e, task vs. control, average of 0.425 vs. 0.405, P = 5.19 × 10^−4^, Mann-Whitney U test) whereas coherence in the broadband 2–40 Hz range (excluding the 5–10 Hz range) was not significantly different between task and control (Fig. S4). This was also true for the 5–10 Hz coherence between the putamen and the central ECoG (approximately above posterior primary motor cortex, task vs. control, average of 0.379 vs. 0.364, P = 3.56 × 10^−11^, Mann-Whitney U test).

We then proceeded to detect CSSC events in the same manner as we did for rodents, based on a putamen 5–10 Hz power threshold as well as a threshold for coherence between putamen LFP and the frontal ECoG. We found that task performance was associated with an increase in CSSC event density (Fig. 5f, task vs. control, 4.70 events/min vs. 3.30 events/min, P = 2.35 × 10^−5^, Mann-Whitney U test), duration (Fig. 5g, task vs. control, 1.68 s vs. 1.38 s, P = 1.13 × 10^−6^, Mann-Whitney U test) and overall probability during NREMS (Fig. 5h, task vs. control, 9.36% vs. 6.05%, P = 1.57 × 10^−5^, Mann-Whitney U test). Together, this suggests that CSSC can also be found in NHPs, and that it may represent a cross-species sleep oscillation that is poised to drive memory consolidation in the cortico-striatal system.

## Discussion

Our results identify a coherent 5–10 Hz oscillation in M1 and DLS during NREM sleep that drives kinematic refinement, likely by coordinating cross-regional reactivation of waking experience. Such cortico-striatal sleep coupling (CSSC) events are mechanistically poised to drive M1–DLS cross-area ensemble binding and memory consolidation by entraining cortical ensembles which were coactive during preceding waking behavior, thereby effectively recruiting downstream DLS neurons^13–16,43,44^. This process seems more effective in potentiating behaviorally significant M1– DLS ensembles. This might rely on “tagging” of cortical or cortico-striatal synapses through neuromodulation (NE^45^, DA^46,47^) during wakeful behavior. Our finding that CSSC activity becomes stronger during mid-late learning is likely related to previous works reporting cortical motor map reorganization, synaptogenesis and potentiation of dendritic spines only after several days of skilled motor learning^48,49^.

Our results also demonstrate that CSSC relies on NMDA receptor activation in DLS. In vitro and in vivo work has shown that NMDA receptors may be important in maintaining striatal “up-states” driven by converging cortical input^28,29^. These transient depolarizations could favor medium spiny neuron activation to facilitate plasticity. During mid-late learning, the increase in cortical entrainment to the 5–10 Hz oscillation might support the ensemble activation needed to elicit downstream striatal “up-states”.

It is likely that sleep related oscillatory dynamics may also have implications for M1–DLS coupling during task performance. For example, past work has found increases in M1–DLS spiking coherence during wake performance between early and late stages of learning a more abstract neuroprosthetic skill involving volitional control of ensembles^50,51^. Interestingly, animals with a local NMDA manipulation did not either learn or develop such coherence. Similarly, we previously found that training across days on a reach-to-grasp task was associated with changes in M1–DLS coherence during awake task performance that emerged over 24-hour periods and could be detected during offline states, including NREMS^13,17^; however, we did not recognize these increases as episodic oscillatory events nor causally link them to refinement of kinematics.

CSSC and sleep spindles appear to play complementary, but dissociable, roles in sleep-dependent learning. Consistent with a large body of literature, early in learning, when rapid improvements in success rate dominate, sleep spindles are elevated and may support cortico-cortical processing, perhaps consistent with their well-established role in early-stage memory consolidation^26,27,52^. Notably, spindles are classically thalamic in origin, reflecting thalamo-cortical coordination^26^. While they are differentiated into fast and slow spindles in humans, work in rodents may not support such a distinction^26^, and rather suggests that they may be grouped into anterior, posterior and global spindles. Anterior spindles, spanning the cortical areas studies here, exhibited a mean frequency of 11.7 Hz^53^. In contrast, the later phase of learning, characterized by kinematic refinement and strengthened cortico-striatal interactions^13–16^, is marked by the emergence of CSSC, a 5–10 Hz oscillation coherent between M1 and DLS. Unlike spindles, CSSC is anatomically and mechanistically tied to cortico-striatal circuits, and our NMDA receptor blockade experiments in the striatum demonstrate that its expression depends on intact striatal processing. Functionally, CSSC drives repeated, temporally precise cross-region reactivation of task-specific activity, suggesting a role in reinforcing cortico-striatal synaptic plasticity and coordinating manifold-level activity across regions to support the transition to automatic behavior. It is possible that while spindles may preferentially support cortico-cortical and cognitive aspects of early learning, CSSC in the DLS may be specialized for motor skill refinement through striatum-dependent cortico-striatal coupling during prolonged training. Notably, aside from sensorimotor functions, cortico-striatal projections are known to support associative and limbic processing^54^. Future work will be needed to study the role of CSSC events during sleep in other types of learning subserved by the striatum.

Notably, consistent with prior work^20^, we observe that M1–mPFC coupling increases early in learning, a phase marked by rapid gains in accuracy and stabilization of task performance, and remains elevated into later stages, suggesting a role for prefrontal–motor interactions in establishing and maintaining task structure and goal-directed representations. CSSC also emerges later in learning as behavior becomes more automatic, selectively coupling M1 and DLS and supporting kinematic refinement. Importantly, our data suggest that the increase in M1– mPFC coupling precedes the slower emergence of CSSC, raising the possibility that mPFC–M1 interactions dominate earlier, cognitively demanding phases of learning, whereas CSSC reflects a later, striatum-dependent mechanism that reinforces task-specific population dynamics within cortico-striatal circuits. Future work will be needed to determine how slow oscillation–mediated activity in mPFC interacts with CSSC, and whether these processes operate sequentially or in a coordinated manner to support different stages of learning.

In mice, CSSC emerges gradually, becoming prominent only during later stages of learning when task accuracy stabilizes and slow kinematic refinement occurs, suggesting a role in consolidating behavior once a reliable cognitive model has formed. In NHPs, we observed a similar cortico-striatal rhythm in putamen and frontal cortex during sleep following task performance. However, unlike mice undergoing new learning, NHPs expressed CSSC after a single day of learning new contingencies. This difference may reflect the presence of an established cognitive schema in primates, built through extensive pre-training, upon which new associations are rapidly incorporated. In contrast, rodents must first acquire both the task structure and sensorimotor mappings, delaying the emergence of CSSC. Thus, across species, we propose that CSSC appears when new information becomes sufficiently structured and predictive to be integrated into existing cortico-striatal networks. This framework may align with human learning, where rapid acquisition often builds on prior knowledge. We thus predict that CSSC and automaticity may appear much faster in primates, especially when it is built on existing knowledge.

In summary, our results indicate that CSSC is important for striatum-dependent consolidation. Such progressive consolidation ultimately leads to automaticity, which can come at the expense of behavioral flexibility^12,18,55^. Because heightened automaticity and the resulting states of behavioral inflexibility are deeply implicated across a spectrum of neurological and psychiatric conditions, mapping how these sleep-dependent networks stabilize or over-consolidate neural pathways offers a new framework for understanding and treating disorders defined by circuit-level rigidity, such as addiction, depression, and obsessive-compulsive disorder.

## Experimental Methods

### Animals and surgery

All procedures were in accordance with protocols approved by the Institutional Animal Care and Use Committee at the San Francisco Veterans Affairs Medical Center. Male and female 3 to 9 months old C57BL/6J mice (C57BL/6J, JAX #000664) were used in this study. Animals were kept under controlled temperature and a 12-h light/12-h dark cycle (lights on at 6:00 AM).

All surgical procedures were performed using sterile techniques under 1.5–5% isoflurane anesthesia. Local treatment with bupivacaine was given before any incision. In all animals, a titanium headbar was implanted to allow head fixation for sleep and task recording. A reinforcement screw was added above the right hemisphere for structural stability. Metabond and dental cement were used to affix implanted elements to the skull. In animals used for Neuropixels recording, the headbar was combined with a well allowing access to the M1–DLS and mPFC probe insertion coordinates (caudal and rostral forelimb areas at AP +0.5 mm, ML 1.5 mm and AP +1.9 mm and ML 1.1 mm, respectively, all from bregma), in the left hemisphere. After habituation and training, these animals underwent a craniotomy and durectomy in preparation for acute recording. For optogenetic experiments, animals were first injected with AAV5-hSyn-eNpHR3.0-EYFP (Addgene #26972) in the DLS (AP +0.5 mm, ML 2.2 mm from bregma, DV 2.2 mm from brain surface). Two weeks thereafter, a single surgery was performed where a headbar was implanted along with a left hemisphere DLS optrode consisting of a neural probe (32-channel 35 μm tungsten microwire electrode array, Innovative Neurophysiology) and a fiberoptic cannula (200 μm, numerical aperture 0.37, tapered tip, Doric. Same coordinates as for injection above). In this surgery, a ground screw was implanted above the left occipital area, connected to the optrode through a silver wire and embedded within the dental cement. For the acute Neuropixels recording, channel location was histologically confirmed by coating the probe with a fluorescent dye (Vybrant DiI cell-labeling solution, Invitrogen) prior to insertion, and tracing the trajectory after sacrifice. Postoperatively, the animals were treated with 0.078 mg buprenorphine. All animals recovered for 7 days before the start of the behavioral procedures.

### Sleep and behavioral training

Seven days after surgery, mice were handled for 5 minutes and were introduced to the automated behavioral box without head fixation. Animals were then habituated to head fixation through daily bouts of increasing duration: 10 minutes, 20, 40, 60, 90, 120 and 180 minutes. Training sessions started at 8:30 AM to match the innate daily sleep timing. Mice were habituated to sleep head fixed, and when they reached sufficient sleep efficiency (usually within 7 days) they were started on a water scheduling regime, where they received a total of 1 mL of water daily. Once water-scheduled, animals were trained to reach for a water drop. They were first allowed to drink directly from the spout delivering the drop. Then the spout was gradually moved back to its final designated position and the animals were whisker-stimulated to reach for newly introduced drops (no auditory cue) with their right forelimb. Once the animals reached a criterion of 5–10 self-initiated reaches to the drop, the training concluded and craniotomy was performed. Throughout reach training, animals continued to sleep head fixed after the task session.

Following craniotomy, the animals were started on a six-day head-fixed task and sleep recording regime. Each day, animals slept for one hour, then performed the task (usually 100–150 trials, 30–45 minutes in total) and then slept for some additional 1.5 hours. Of the 6 animals used for electrophysiology, one animal had only behavior data for day 6 (and behavior and electrophysiology as usual for days 1-5).

All elements of the behavioral task were controlled by a Raspberry Pi single-board computer and custom Python scripts. Each trial started with a 2-s enforced hold period where animals had to keep their forepaws on two plastic posts. Once an automated video script detected one of the paws leaving the immediate surroundings of the posts, the 2-s count would restart. Following this 2-s period, a 4 kHz auditory cue would sound, an 8 microliter drop would be dispensed from the spout and the animal would be given 7 s to reach. After 7 s, a servo motor would remove the drop in preparation for the next trial. After a 5-s inter-trial period (including drop removal), animals would be allowed to start the 2-s hold period for the next trial. Animals were exposed to the trial structure and to its different elements only on day 1 of the recording.

All trials were captured by two video cameras (Basler, 150 frames/s) placed on the side and front of the behavioral box. Frame grabbing was synchronized with the electrophysiology data using a Raspberry Pi digital output.

### Behavioral analysis

Behavioral analysis relied on an automated tracking algorithm (Deeplabcut^56^) for body parts (hand, fingers, tongue) and task-relevant components (spout, drop, plastic posts). All behavioral events (e.g., trial start, reach onset, reach end) and measures (e.g., reaction time, trial classification, reach trajectory) were extracted from Deeplabcut tracking data automatically by a custom script. Trial start was defined as the point in time where the drop was introduced. Reach onsets and ends were detected as follows: a reach threshold was defined as the first timepoint, after trial start, when the hand transversed a certain percentage of the distance from its initial position to the spout. Then, reach onset was defined as the last timepoint, before reach threshold, where the hand velocity crossed a predefined velocity threshold. Reach end was detected as the first peak in the hand distance from its initial position, after reach threshold. Subsequent reaches were detected similarly after the first reach end. Velocity, distance and duration criteria were similar between animals and were constant between days within the same animal.

Reaction time was defined as the latency from trial start to reach onset. A trial could either be successful (animal reached and successfully grabbed the drop within 1 s after the end of the first reach), missed (animal reached, but the drop was missed, knocked off or not grabbed within 1 s after the end of the first reach) or omitted (animal did not perform any reaches during the trial). Success rate was quantified as the number of successful trials out of all non-omitted trials. For reach trajectory and reach endpoint variability, we only used the first reach in every non-omitted trial. Trajectories were given in x and y coordinates from the side camera (anterior-posterior and inferior-superior, respectively). Reach trajectory distance was quantified per day as the Euclidean distance of the trajectory of a given reach, from the average trajectory across all reaches on that day, averaged across the x and y dimensions. Seeing as different reaches had slightly varying durations, whereas Euclidean distance is calculated between same-length vectors, all reaches were interpolated to a common duration, defined as the 75^th^ percentile of all reach durations in that session. Similarly, the endpoint variability was quantified as the 2D standard deviation of the distribution of reach endpoints (defined as the mean of the last 10% of frames before reach end, separately in x and y coordinates) relative to the daily mean of the reach endpoints. For three task sessions, of all animals and days, one camera was losing frames such that behavioral analysis could not rely on the data it generated. In these cases, we performed manual classification of trials including detection of trial starts and reach onsets and ends based on the unaffected front camera. To compare endpoint and trajectory variability across all days, side camera endpoint and trajectory variability values were estimated based on those obtained from the front camera, using linear regression relying on the remaining sessions from the same animal, where both the frontal and side camera produced videos with an equal and full number of frames.

### In-vivo electrophysiology

For acute electrophysiological recordings, spiking activity and LFP were recorded using two Neuropixels 1.0 probes (imec). Spiking data was sampled at 30,000 samples/s and filtered at 300–6000 Hz, and LFP data was sampled at 2,500 samples/s and filtered at 0.5–300 Hz. Spike sorting was performed offline using Kilosort 4.0. Curation was first done automatically (using bombcell) and then manually (using Phy): All units were visually evaluated to ascertain proper signal-to-noise ratio and waveform amplitude, clear cluster boundaries and that the vast majority of detected spikes had ISI > 2 ms. UnitMatch^57^ was used for identification of units recorded during task sessions and subsequent sleep sessions. For optrode recording, data was acquired using an RZ2 system (Tucker-Davis Technologies), where data (spiking and LFP) was sampled at 24,414 samples/s. Spike sorting was performed offline using Mountainsort 5.

### DLS infusions

To test whether cortico-striatal 5–10 Hz activity relies on striatal NMDA receptor activity, we infused 400 nL of either saline or the NMDA receptor antagonist AP-5 (D-(-)-2-Amino-5-phosphonopentanoic Acid, Sigma-Aldrich 165304, 5 µg/µl, infusion rate 100 nL/min) into the left DLS (AP 0.5, ML 2.2 from bregma, DV 2.2 mm from brain surface) immediately following task training, in 4 animals for 6–8 days. Immediately after infusion, a single Neuropixels 1.0 probe was inserted to record M1 and DLS activity, and the animals were allowed to sleep. Saline vs. AP-5 days were randomized.

### Behavioral state classification

Behavioral states (NREM sleep and wakefulness) were classified based on cortical (M1) LFP and movement data extracted from video taken at 50 frames/s during pre-and post-task sleep sessions. Head fixation allowed high resolution video from a close range where minimal movement was readily registered.

LFP was preprocessed by manual artifact rejection and rejection of noisy channels, if present. Then, it was z-scored across the entire sleep session recording period and the cortical mean LFP activity was extracted. Video and LFP data were then segmented into non-overlapping 5-s epochs which were individually staged as wakefulness, NREM sleep or unclassified, based on the following procedure^17,20^: For each 5-s epoch, an LFP high-to-low log power ratio was calculated as the ratio between LFP power at 30–55 Hz and power at 0.5–4 Hz. The degree of movement was quantified as the mean absolute change in pixel intensity between consecutive frames, subsampled at 2 Hz, in an ROI of the frame that includes only the animal.

A k-means classifier was then used to classify all epochs into two clusters, based on detected movement and LFP power ratio. This procedure generated a cluster with low movement and low log LFP power ratio (i.e. more 0.5–4 Hz activity, “putative NREMS” cluster) and a cluster with higher movement and higher ratio. Within the putative NREMS cluster, only epochs with zero or minimal movement were considered true NREMS epochs, and all the rest were considered unclassified. All single NREMS epochs that were flanked by wake on both sides were excluded from analysis.

### NREM sleep rhythm detection

Detection was performed separately for M1 and DLS, and separately per-channel in each region. M1 usually spanned 110–150 channels (corresponding to 1.1–1.5 mm), and DLS usually 190– 250 channels. Specific region boundaries were based on reconstructed probe trajectories and neuronal densities post spike detection and curation. For LFP analysis, 20 channels were analyzed for each area, spanning its depth. DLS data was median-centered to reduce possible effects of LFP volume conductance from cortex.

Detection of slow oscillations (SOs) and sleep spindles relied on conventional detection algorithms^17,35^. Briefly, consecutive NREMS epochs were analyzed as sleep bouts. For SOs, LFP data from each channel was filtered at 0.5–4 Hz (6th order zero-phase Butterworth high-pass filter followed by a 10th order zero-phase Butterworth low-pass filter). Positive-to-negative zero-crossings were then identified along with the preceding peaks, consecutive troughs and surrounding negative-to-positive zero crossings. Candidate events were classified according to the following SO criteria: 1) Peak in top 15% of peaks, 2) Trough in top 40% of troughs (i.e., most negative), and 3) Latency between the negative to positive zero-crossings between 150 and 500 ms.

For sleep spindles, the LFP data was first filtered at 10–16 Hz (8th order zero-phase Butterworth filter). The instantaneous amplitude of the signal was then obtained by applying the Hilbert transformation and smoothed by a 200 ms Gaussian kernel. The distribution of smoothed spindle range Hilbert amplitudes across all NREMS for a given channel was obtained. Two thresholds were used to define sleep spindle events: the mean plus 2 standard deviations of the above distribution (high threshold), and the mean plus one standard deviation (low threshold). Putative events exceeding the high threshold for at least one sample, exceeding the low threshold for at least 500 ms, and overall shorter than 2 s, were defined as sleep spindles. Spindles that were <300 ms apart were joined into one event. The use of 2.5 and 1.5 standard deviations above the mean instead of 2 and 1, respectively, returned similar results for all analyses relying on sleep spindle detection.

### CSSC event detection

Cortico-striatal sleep coupling (CSSC) events were detected separately for each of the 20 analyzed channels in M1 and DLS. A detected CSSC event had to (1) be at least 1 second long, exhibit (2) 5–10 Hz power higher than the 70th percentile calculated across all NREMS for that channel, and exhibit (3) M1–DLS coherence of 0.7 or higher. Below is a detailed description of the detection algorithm, with justification for the selected thresholds.

First, all NREMS epochs were segmented into one-second segments. For each channel, the mean 5–10 Hz instantaneous amplitude of each segment was derived by first filtering the signal at 5–10 Hz (6th order zero-phase Butterworth filter) and then obtaining the instantaneous amplitude using the Hilbert transform of the filtered signal. Only 1-s segments in the top 30% across NREMS were considered for further analysis. This threshold was chosen as traces showing above-threshold 5–10 Hz exhibited a strong oscillation that was discernible by inspection and detectable as a prominent peak in the power spectrum.

The next stage involved calculating the coherence associated with each qualifying 1-s segment. To do so, we defined five channels in M1 and five channels in DLS which were used for calculating the coherence with all analyzed DLS and M1 channels, respectively. The M1 channels were chosen to span the 0.25 mm just ventral to the center of M1. The DLS channels were chosen to span the 0.25 mm just dorsal to the center of DLS. These were chosen due to their central positions within M1 and DLS. M1–DLS 5–10 Hz coherence was then calculated for each M1 channel with the five DLS channels separately, and the average coherence across the five pairs was obtained. One-second segments which exhibited both high M1 5–10 Hz power and M1–DLS coherence higher than 0.7 were defined as M1 CSSC events, and similarly for DLS (i.e., high DLS 5–10 Hz power and high M1–DLS coherence, calculated between the DLS channel in question and each of the five M1 channels).

The choice of 1-s segments was motivated by the fact that coherence estimation relies on relatively long traces of data to be accurate. The choice of the 0.7 coherence threshold was based on our analysis of the distribution of coherence values between all channels in M1 and the five DLS channels above. We fit a Gaussian mixture model to the pooled distribution for 5–10 Hz coherence values across all sleep sessions along the learning trajectory and all animals. Then, we used the Bayesian Information Criterion (BIC) to deduce the number of underlying Gaussians that best fits the distribution. The BIC showed considerable improvement for a mixture of four Gaussians, then plateaued. The transition point between the top 5–10 Hz M1–DLS coherence Gaussian and the lower ones was found to be at 0.689, and a threshold value of 0.7 was used to reduce type I errors.

Notably, post-hoc detection of CSSC events in animals from the optogenetic intervention cohort relied on a slightly modified approach. We detected CSSC events as segments at least 1-second long in the top 30% of DLS 5–10 Hz power, i.e., the power and duration requirements were identical to the ones used for the acute recording animals. However, M1–DLS coherence was not considered as for these animals we recorded only DLS and not M1. To reflect the difference, we refer to these CSSC events as “putative CSSC events” in the text.

### Spiking and neuronal coupling analysis

Spiking analysis was performed only on well-isolated curated neurons in all channels of the region at hand, provided they were recorded for at least 85% of the duration of the session. We calculated the pairwise coupling between two simultaneously recorded M1 and DLS neurons, during either NREMS or task performance. We first found the number of simultaneous spikes per pair as the number of spikes from neuron A that had a spike from neuron B less than 40 ms away. We then z-transformed this number relative to the mean and standard deviation obtained from a distribution of jittered spikes for one neuron relative to the other (100 jitters per pair). A pair was considered coupled if its z-score was larger than 3. To compare coupling during NREMS, we separately performed this process for spiking of the same pair during CSSC events and outside of CSSC events (1-s NREMS segments exhibiting bottom 30% of 5–10 Hz coherence, to allow comparison of a similar duration of time between groups). A pair was considered coupled if its average z-score across the two conditions was > 3. To control for an increase in coupling that reflects a state of generally elevated M1–DLS coherence, rather than a specific increase in the 5–10 Hz band, we performed the same analysis on coupling during CSSC vs. during periods of high broadband coherence (segments exhibiting top 30% of 2–40 Hz coherence excluding the 5– 10 Hz band, Fig. 2). A latency of 40 ms is higher than expected for single synapse transmission latency between M1 and DLS. We chose this longer latency to detect the cofiring of ensembles of M1 and DLS neurons which may not all necessarily be separated by only one synapse.

### Manifold dynamics: decoders and reactivation analysis

#### Decoding of DLS trajectories from M1 spiking (Fig. 3)

We used Gaussian Process Factor Analysis (GPFA^33^) to obtain low-dimensional representations of reaching- or resting-related neural activity during waking task performance. For each session, we concatenated all single-trial single-neuron binned spike counts (from 4 s before to 4 s after reach onset, 40 ms bins) and z-transformed the result to account for differences in mean firing rates. GPFA was then run on the neuron × bins matrices for all neurons recorded in a single session. Single factor trajectories were obtained for the top 3 factors independently for DLS. To explore changes in manifold-level coupling between M1 and DLS with learning, we utilized a decoding approach where M1 spiking was used to predict simultaneously recorded DLS GPFA neural trajectories. A ridge-regularized linear regression model was trained on the cross-trial concatenated data (0.24 s before to 0.16 s after reach onset) using 5-fold cross-validation. To decode DLS GPFA trajectories, the decoder could rely on the current M1 activity bin, as well as on the next and previous single bins. Notably, GPFA calculation was performed on a wider time window (±4 s around reach onset) than model training (0.24 s before to 0.16 s after reach onset) to ensure that the resulting trajectories capture the full scope of neural dynamics across the task. For each session, we calculated Pearson’s correlation between the predicted and actual neural trajectories per GPFA factor. To avoid non-learning related effects on decoding accuracy, we used a constant number of trials and numbers of M1 and DLS neurons across days. In sessions with more than the animal-wise minimal numbers of trials or neurons, we used batches to account for all available neurons and trials and averaged all per-batch correlation values. For non-reach controls, we used same-duration (0.4 s) segments spanning the middle of the wait period before trial start after verifying it was not associated with apparent movement. For a single animal, days 3 and 6 were excluded from this analysis due to low neuronal yields (day 3) or lack of electrophysiology recording (day 6, therefore Fig. 3e has 28 points instead of 30).

#### Decoding of DLS trajectories from M1 trajectories during NREMS (Fig. 2)

A similar GPFA decoding approach was used to decode DLS neural trajectories based on M1 trajectories strictly during NREMS. Here, we used 1-s NREMS segments, where each segment was associated with a specific M1–DLS LFP 5–10 Hz coherence, and the individual GPFA trajectories in M1 and DLS (25 bins of 40 ms). A similar ridge-regularized linear regression model was used here to decode per-factor DLS activity based on the top 3 M1 factors, and to relate the decoding accuracy (reflected as the per-second correlation between actual and predicted DLS activity) to M1–DLS 5–10 Hz coherence.

#### Reactivations of task GPFA-derived factor during sleep (Fig. 2)

We used GPFA neural trajectories to monitor reactivations of reaching- and resting-related information during NREMS. Binned NREMS spiking activity for the same neurons recorded during task performance (neurons recorded across sessions were detected using UnitMatch) were projected onto the task-derived factors, to obtain single-factor single-region reactivation profiles. To obtain per-bin cross-factor reactivation strength for M1 and DLS, we averaged the absolute reactivation strengths over the top four factors, independently in M1 and DLS. Joint reactivations were defined as simultaneous reactivation events in both M1 and DLS (i.e. occurring in the same 50-ms time bin), exceeding the 80% percentile of their respective distributions of reactivation strength peaks. For CSSC and sleep spindle detection related to reactivation analysis, we used one central M1 or DLS channel for detection as event timings slightly varied between channels. For both M1 and DLS, we used the middle channel of the five channels used for M1 and DLS coherence calculation (see under CSSC event detection above). Reactivation analysis relied on the registration of the same M1 and DLS units throughout both task and sleep sessions. Of the 36 pairs of task and sleep sessions (6 pairs across 6 animals), 2 had low cross-session neuronal yields (on day 3 and day 1) and in one electrophysiology was not recorded (day 6). Thus, eleven sessions were analyzed for each learning stage — early, mid, and late — for a total of 33 sessions. For the PLV of reactivation events during CSSC analysis (Fig. 2h), only sessions with more than 110 reactivation events during CSSC were considered for PLV calculation, to avoid spuriously high PLVs.

### Optogenetic manipulations

Wild-type mice (C57BL/6J, JAX #000664) were injected with a viral vector carrying the inhibitory halorhodopsin (AAV5-hSyn-eNpHR 3.0-EYFP (Addgene #26972) into the left DLS (AP 0.5, ML 2.2 relative to bregma, DV 2.2 mm from brain surface). After the implantation of an optrode (cannula + microwire array, see above) the animals were habituated for sleep and trained for the task, as for the acute recording cohort. For the optogenetic inhibition experiments, animals learned the task for 11 days, instead of the usual 6 days, and optogenetic intervention was performed during the sleep session immediately after task performance. On days 1–5, animals were subjected to a 5–10 Hz oscillation inhibition regime, and on days 6–10, they were subjected to a matched-duration random timing inhibition regime, where the same overall per-session duration of inhibition and the same average individual laser ON time were used (as in the 5–10 Hz inhibition), but randomly throughout sleep. For behavioral analysis, days 2–6 were compared to days 7–11 as within-animal inhibition vs. matched random interventions. For the 5–10 Hz oscillation inhibition, DLS LFP was used to detect online increases in oscillation power. When the RMS of the 5–10 Hz filtered signal (averaged on 6 DLS channels) exceeded a pre-defined cross-session threshold for a duration longer then 0.3–0.4 s, the laser was activated until the RMS decreased below the above threshold (or a maximum of 3 s). We used a 532-nm laser (Opto Engine LLC) and the power measured at the tip before each session was set at 15 mW. After the final 5–10 Hz oscillation inhibition session on day 5, the average per-session overall stimulation duration as well as the average individual laser ON time were obtained and used for matched random inhibition on days 6–10. For both regimes, inhibition was only used when the animal was in apparent NREM sleep (no movement in the video, and slow wave or 5–10 Hz activity in the LFP). Black heat-shrink tubing was used to prevent laser illumination from being detected by the animal and affect sleep.

### NHP recordings

The task scheme and sleep and recording procedures are reported in our previous work^58–60,39^. Briefly, data was obtained from two young adult, female vervet monkeys (Chlorocebus aethiops, D and N) weighing ∼3.5 kg. The animals were trained for a period of 3–4 months on a classical conditioning task where they would be headfixed in front of a screen presenting fractal cues (adapted from www.easyfractalgenerator.com). Each trial started with the presentation of a cue for 2 s, followed by the delivery of either a positive (juice reward), negative (airpuff to the eye) or neutral outcome (no outcome). A 5–7 s inter-trial interval (ITI) separated trials. Ten such rewarding, aversive and neutral trials were randomly interleaved to create a single 30-trial block. Trial blocks could be either overtrained (i.e., the animal knew the cues and their associated outcomes from previous training) or novel (i.e., the cues were never presented to the animal before and the animal could not predict the outcome). Also, trial blocks could either be deterministic (where positive and negative outcomes were always delivered) or probabilistic (where outcome delivery probability was 0.5 (animal D) or 0.75 (animal N)). In total, animals usually performed 4 block sessions prior to sleep, including one of each block above (overtrained deterministic, overtrained probabilistic, novel deterministic and novel probabilistic, in random order). The animals were able to learn the novel contingencies well and performed preemptive licking (for rewarding trials) and blinking (for aversive trials) after several trials^39^.

The animals were habituated every night for 2–3 months to sleep head fixed in a primate chair, positioned in a dark, double-walled sound-attenuating room. This limited head movement but otherwise allowed the animals to sleep in a position similar to their natural sitting sleeping posture. At the end of habituation, the animals slept in the room the entire night (10-11 PM until 5-6 AM, 4-5 nights a week). After the habituation, the animals displayed normal sleeping patterns and high sleep efficiency^59^. Polysomnographic recordings were used for behavioral state classification. We acquired ECoG and EMG signals, alongside continuous infrared video (at 50 frames/s) to determine whether the eyes were open or closed. ECoG activity was collected from five epidural screws spanning frontal (F_3_), central (C_1_, C_4_), and posterior-occipital (PO_3_ and PO_4_) regions, using the standardized 10–20 system. ECoG channels were referenced to two titanium epidural ground screws that were interconnected and positioned at P_5_ and FC_6_. Muscle activity was recorded from the right trapezius using bipolar needle electrodes selected for their strong and discriminative signal across sleep states. ECoG was sampled at 2,750 Hz and bandpass-filtered at 0.1–35 Hz (zero-phase Butterworth filter, stopband at 0 to 0.05 Hz, 40 to 1,375 Hz) and EMG was similarly filtered at 10–100 Hz (stopband at 0 to 5 Hz, 120 to 1,375 Hz). Positioning of electrodes, sampling, and filtration followed the recommendations of the American Association of Sleep Medicine (AASM) Manual for the Scoring of Sleep and Associated Events.

Sleep stages were identified using a semiautomated approach applied to 10 s non-overlapping epochs, combining three features: the ratio of high- to low-frequency ECoG power (average 15– 25 Hz power divided by average 0.1–7 Hz power), EMG signal RMS, and the proportion of time the eyes were open (derived from video-based pixel analysis). These features formed clusters corresponding to wakefulness (high EMG RMS, increased EEG high/low ratio, eye-open fraction close to 1), NREMS (low EMG RMS, decreased EEG high/low ratio, eye-open fraction close to 0), and REMS (very low EMG RMS, increased EEG high/low ratio, eye-open fraction close to 0), which were validated against expert scoring, requiring high agreement to proceed.

Daily recording sessions included task performance and the subsequent full night’s sleep. To be compatible with the rodent sleep recording durations, we only analyzed NHP data from the first hour of sleep. Electrophysiological data was collected using eight glass-coated tungsten electrodes which were advanced separately toward the putamen. Electrical activity was filtered using a hardware Butterworth filter at 0.075–10,000 Hz and sampled at 44 kHz. Spiking activity was sorted online using a template matching algorithm (SnR; Alpha Omega Engineering). The putamen was identified according to its stereotaxic coordinates, based on MRI and primate atlas data and real-time assessment of the electrophysiological features of neurons.

Overall, 18 sleep sessions from two animals were analyzed (D, 14 sessions (7 task, 7 control), N, 4 sessions (3 task, 1 control)). Sleep spindle and CSSC event detection in the NHP data followed the same procedures used for mice and outlined above. The only changes made for the NHP were the use of 2.5 and 1.5 standard deviations above the mean (NHPs) instead of 2 and 1 (mice) for spindle detection and utilizing a top 40% 5–10 Hz power criterion (NHPs) instead of 30% (mice) for the CSSC event detection. Additionally, the frontal ECoG–putamen coherence threshold was slightly modified between the two animals (0.7 for D, and 0.6 for N. The threshold for animal N was reset to match its slightly left-shifted distribution of frontal ECoG–putamen 5–10 Hz coherence values, probably due to a difference in ECoG electrode positioning). 5–10 Hz coherence and CSSC event detection were performed separately for the putamen–frontal ECoG and the putamen–central ECoG, with similar results, as reported in the results section.

### Quantification and statistical analysis

All data analysis was conducted using Python 3.10 and MATLAB 2024a (MathWorks). Unless indicated otherwise, all figures show mean±s.e.m. Across the manuscript, parametric tests (e.g., paired and non-paired t-tests) were used only after data normality and equality of variance were confirmed via the Shapiro-Wilk test and the Levene test, respectively. Otherwise, non-parametric tests (e.g., Mann-Whitney U test and Wilcoxon signed-rank test) were used. In cases where related comparisons were not consistently showing data normality and equality of variance, we used the non-parametric tests throughout all comparisons for uniformity and rigor. Statistical threshold was set at α=0.05 for all analyses.

---

**Fig. S1.**
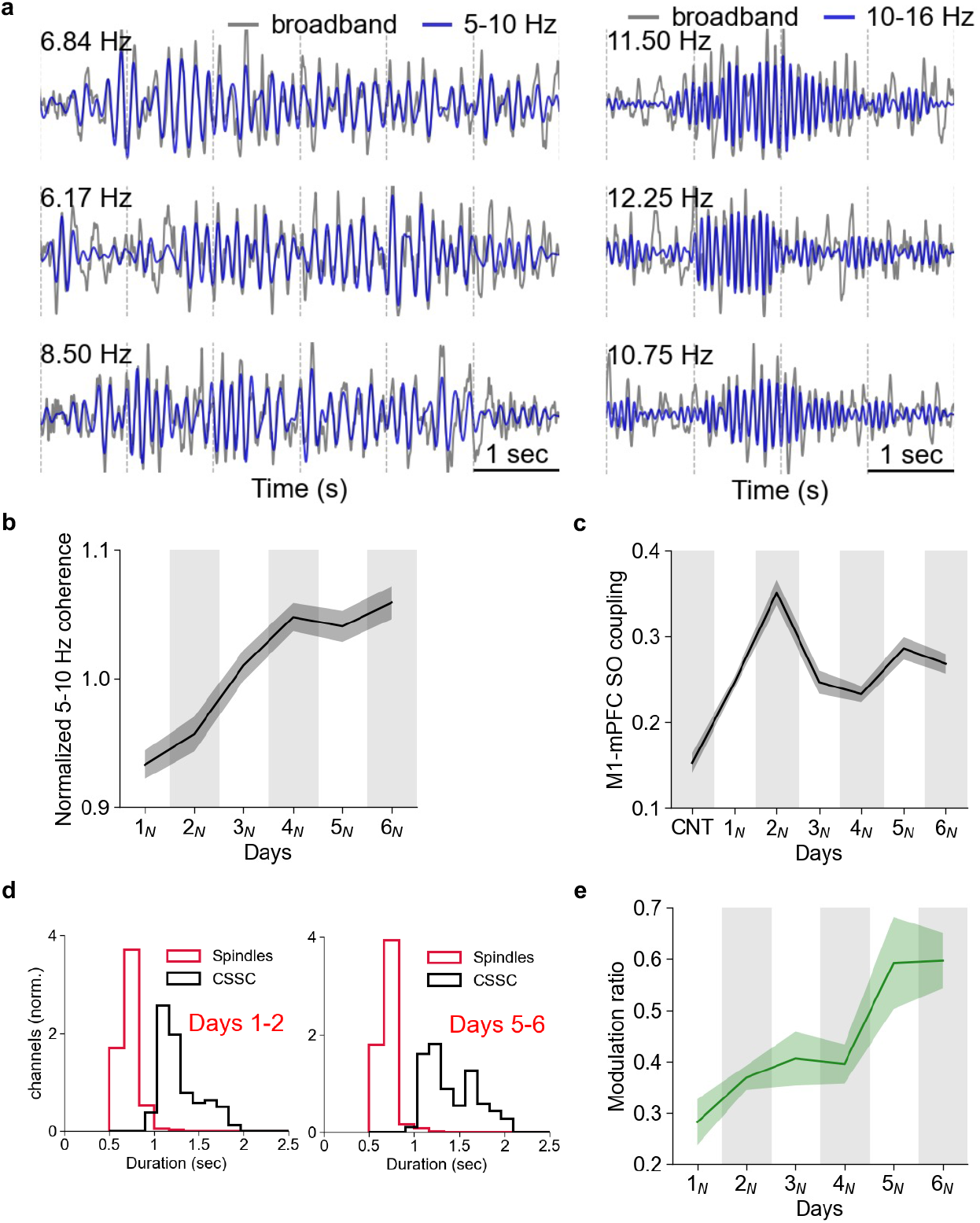
**a,** Left, example DLS LFP traces with representative 5–10 Hz activity from three different animals (days 5, 3 and 5 of learning), distinct from the animal the example in Fig. 1i was derived from. Top left, peak frequency of broadband trace. Vertical lines represent 1-s intervals. Right, example DLS LFP traces with detected sleep spindles from the same animals and using the same scale. Broadband, 2–40 Hz for right and left sides. **b**, M1–DLS 5–10 Hz coherence normalized by the average broadband M1–DLS coherence (2–40 Hz, excluding the 5–10 Hz band) across channels and animals, along learning. **c,** M1–mPFC SO coupling (the proportion of M1 SOs that are 0.2 s or less away from an mPFC SO, out of all M1 SOs) averaged across M1 channels and animals, with learning. Control vs. D2-6 average, 0.15 vs. 0.28, P = 2.45 × 10^−11^, Wilcoxon signed-rank test. **d,** Distribution of all sleep spindle and CSSC event durations (across channels and animals) for D1-2 vs. D5-6. D1–2, spindle vs. CSSC duration, 0.72 vs. 1.29, P = 5.22 × 10^−41^, Wilcoxon signed-rank test. D5–6, spindle vs. CSSC duration, 0.71 vs. 1.43, P = 8.23 × 10^−88^, Wilcoxon signed-rank test. **e,** Modulation ratio along learning, across animals. The modulation ratio is defined per sleep session as the mean M1–DLS 5–10 Hz coherence, divided by the average spindle density in M1 and DLS.

**Fig. S2.**
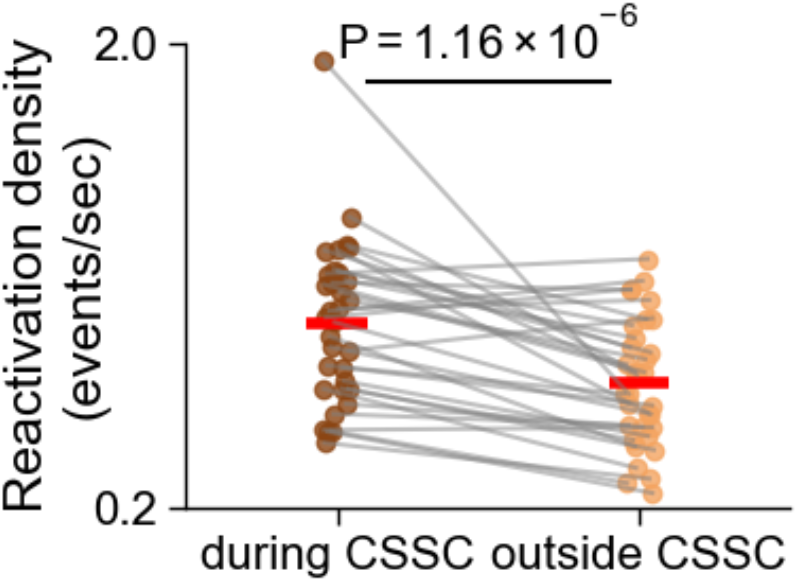
M1–DLS joint reactivation density (number of joint reactivation events per second), during CSSC events vs. outside of CSSC events, across all sleep sessions in all learning stages, 0.916 vs. 0.680 events/sec, P = 1.16 × 10^−6^, Wilcoxon signed-rank test.

**Fig. S3.**
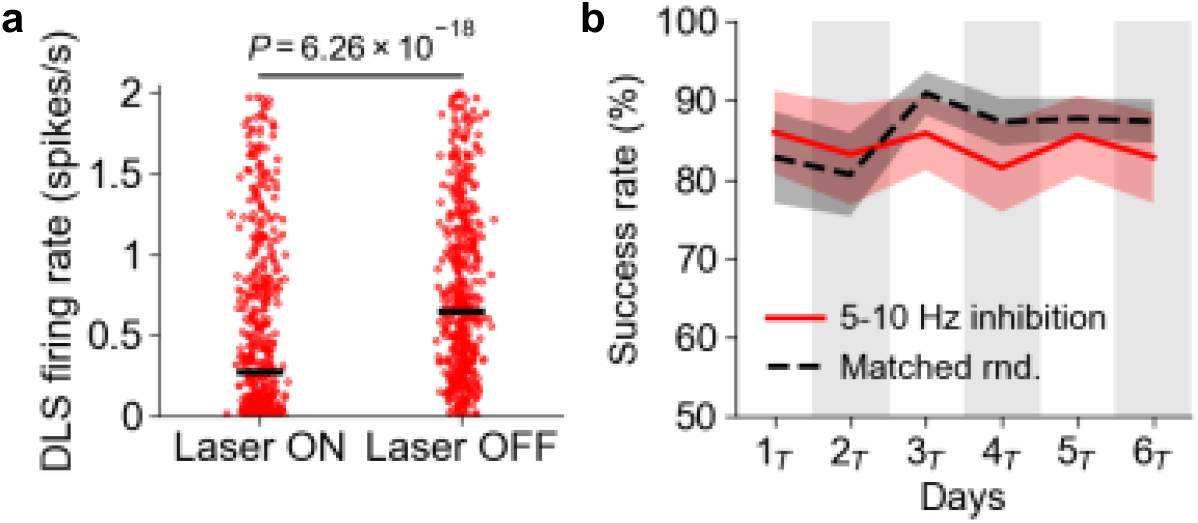
**a,** Firing rates for the same putative medium spiny neurons in DLS (i.e., neurons with average firing rate < 2 spikes/s. n=481 neurons, from all animals and all sessions), during laser ON vs. laser OFF. Median firing rate, 0.27 spikes/s vs. 0.64 spikes/s, P-value, Wilcoxon signed-rank test. Medians are reported instead of means due to the left-skewness of the underlying distribution, especially for laser ON. **b,** Success rate for the 5–10 Hz inhibition and matched random inhibition regimes.

**Fig. S4.**
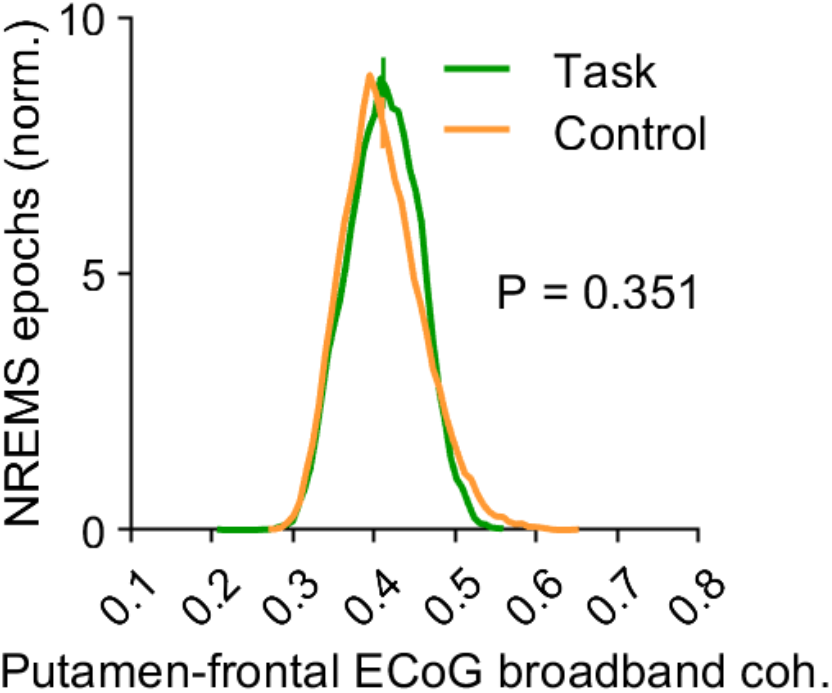
Putamen-frontal ECoG coherence in the 2–40 Hz range (excluding the 5–10 Hz range), in the first hour of sleep after task performance, vs. in the first hour of sleep when no task was performed. Vertical bars, average, 0.411 vs. 0.407, P-value, Mann-Whitney U test

## Notes

### Competing Interest Statement

The authors have declared no competing interest.

### Funding

- NIH: 1RF1NS132913-01
- Veterans Affairs Medical Center San Francisco: 5I01RX001640-12

